# RapidAIM 2.0: a high-throughput assay to study functional response of human gut microbiome to xenobiotics

**DOI:** 10.1101/2022.08.03.502618

**Authors:** Leyuan Li, Janice Mayne, Adrian Beltran, Xu Zhang, Zhibin Ning, Daniel Figeys

**Author notes:** Both authors contributed equally (Leyuan Li, Janice Mayne).

## Abstract

Our gut microbiome functions like an organ, having its own set of functions and roles which can be modulated by various types of xenobiotic and biotic components. High-throughput screening approaches that are established based on *in vitro* or *ex vivo* cell, tissue or organ models greatly accelerate drug discovery and our understanding of biological and pathological processes within these systems. There was a lack of a high-throughput compatible functional screening approach of the gut microbiome until we recently developed the RapidAIM (Rapid Assay of Individual Microbiome). RapidAIM combines an optimized culturing model, which maintains the taxonomic and functional profiles of the human gut microbiome *in vitro*, and a high-throughput metaproteomics workflow to gain deep functional insights into microbiome responses. This protocol describes the most recently optimized 2.0 version of RapidAIM, consisting of extensive details on stool sample collection, biobanking, *in vitro* culturing and stimulation, microbiome sample processing, and metaproteomics measurement and data analysis. To demonstrate the typical outcome of the protocol, we show an example of using RapidAIM 2.0 to evaluate the effect of prebiotic kestose on *ex vivo* individual human gut microbiomes biobanked with five different workflows; we also show that kestose had consistent functional effects across individuals and can be used as positive control in the assay.

## Introduction

Numerous studies have shown that xenobiotic compounds can functionally affect the gut microbiota. These include pharmaceutical compounds, not only those developed for combating microbial infections, but also those non-antimicrobial drugs that were developed to target the host functions[1]. Biotic factors from external sources, such as probiotics, pathogens, phages, etc. also influence the gut microbiome functionality in various ways. To deconvolute the complexity of microbiome responses to these factors, *in vitro* approaches in the absence of the host component have been used. Studies have shown that commensal bacterial species in the gut can be directly affected by 24% of host-targeted drugs[1], and therapeutic drugs can be accumulated in gut bacteria without altering their abundances[2]. However, studying gut microbes in isolation has shown its limitations, since microbial species function differently in complex community settings in comparison to pure cultures. Cooperative and antagonistic interactions collectively contribute to microbiome diversity and resilience[3]. Several synthetic communities have also been used to evaluate the effect of drugs on community composition and cross-feeding interactions[2]. Nevertheless, the natural human gut microbiome harbors hundreds of microbial species, with a considerable variation in taxonomic functions and compositions among different individuals[4–6]. The complexity and variability is much greater than synthetic gut microbial communities. Studies have shown long-term stability of the individual gut microbiome across life span[7], and this stability can be irreversibly perturbed by xenobiotic stimulation[8, 9]. Therefore, when it comes to the context of individual gut microbiomes, it is necessary to adopt an assay that maintains the individuality of the community composition and function *in vitro*. In addition, since evaluating xenobiotic compounds or biotic components against different individual microbiomes require large matrices of samples, high-throughput-compatible models and assay readouts are necessary to perform such studies.

The lack of high-throughput methods to analyze microbiome protein composition limits our understanding of community functional and ecological responses to external perturbation. Fortunately, we previously developed RapidAIM, namely Rapid Assay of Individual Microbiome, taking advantages of fast-pass metaproteomics to study the human gut microbiome responses to xenobiotics in a high-throughput compatible setting[10, 11]. This assay is capable of maintaining stable gut communities *in vitro* in the absence of a host, and has great potential to speed up our acquisition of knowledge of the functional impacts of xenobiotics on gut microbiomes. Liquid chromatography – tandem mass spectrometry (LC-MS/MS) is among the most widely used approach for metaproteomic analysis of microbial community[12]. Briefly, LC-MS/MS separates and analyzes peptides in samples to obtain MS/MS spectra, which are subsequently matched to peptide sequences through database search approaches. With its fast-growing measurement depth and capacity, metaproteomics techniques have used to study microbiome-associated health and disease such as inflammatory bowel disease[13, 14], colorectal cancer[15], diabetes[16], mental illness[17], and COVID-19[18]. It has also been used to evaluate *in vitro* responses of gut microbiomes to various xenobiotics. In RapidAIM 2.0, we incorporate automation and multiplexing technique into the metaproteomic analysis workflow to observe the response of protein expression in the *in vitro* microbiome in a high-throughput manner. The use of isobaric chemical labels, such as tandem mass tag (TMT) approach, offers great potential in the analysis of large sample sets such as those generated from our individual microbiome-xenobiotics assay designs. Tandem mass tag (TMT)-based quantitation has recently been used for large-scale proteomics studies owing to its high multiplexing capacity and deep proteome coverage[19]. The use of TMT labeling also significantly reduce LC-MS/MS time and cost [20]. To enable high-throughput sample analysis, we have developed a streamlined TMT labeling workflow using pre-aliquoted dry TMT in 96-well plates, which achieved comparable labeling efficiency and improved inter-sample consistency for quantitation[21]. This streamlined TMT labeling workflow is fully compatible with automation of the metaproteomics sample preparation steps which altogether can significantly increase the robustness of liquid handling compared to manual operation, speed up the experimental workflow and further increase the throughput[22].

### Development of the protocol

The RapidAIM 2.0 approach is an updated version of the previous RapidAIM series approaches developed and used in research studies by our laboratory [10, 23]. In the previous workflows, we described a 96-well based workflow to study the functional responses of individual gut microbiomes to xenobiotics *in vitro*. The development of the RapidAIM 2.0 protocol consists of three major parts, 1) optimized the culture medium and establishing and validating the 96-well based scalable culturing model[11, 24]; 2) established a 96-well based metaproteomic sample processing and data analysis workflow[10]; and most recently, we 3) developed and validated a live microbiota biobanking workflow that is helpful to increase the reproducibility of experiments[23]. A limitation of the previous protocol was the considerably large sample size and LC-MS/MS time consumption for metaproteomics analysis. Therefore, we updated the protocol to overcome this limitation. New feature and components of the protocol include 1) optimization of the protein extraction and purification protocol; 2) automation of the protein digestion and desalting protocol; 3) introduction of TMT multiplexing technique for labeling and quantitation of peptides and proteins allowing for the analysis of up to 10 samples in one LC-MS/MS run[21]; and 4) a TMT-based statistical analysis streamline for clustering functional responses. We estimate that, for an experiment containing 320 samples (four 96-well plates), this updated workflow requires only six days for culturing and sample processing, and it shortens LC-MS/MS sample analysis time from approximately 20 days to 3 days.

### Applications of the method

RapidAIM has been widely used in numerous research studies. As a proof of concept of RapidAIM, we first used it to assess the effect of 43 xenobiotic compounds on five individual gut microbiomes and discovered that seven of the tested compounds showed consistent effects across individual samples, while some other compounds showed significant but individually distinct effects[10]. RapidAIM was then used to evaluate the effect of a panel of structurally similar compounds of berberine[25], different structures of resistant starches[26] as well as commonly used sweeteners[27] on individual gut microbiomes. RapidAIM was also used to evaluate the effect of a bacteriophage preparation on microbiome composition and function[28], as well as being applied in two ongoing clinical trials for the selection of therapeutic interventions (NCT04520594 and NCT04522271). Following our RapidAIM publications, we’ve received many inquiries about experimental details of the protocols, as well as collaborations and interests from industrial and clinical partners. Our optimized culture model of RapidAIM has been widely used by other researchers to investigate gut microbiome responses to bile[29], oligomannate[30], dietary fibers[31], antibiotic[32], and nanoparticles[33] etc. With the rapid rise of the automation and big data era, our development of the 2.0 version of the protocol to expand the capability to study microbiome responses to various stimuli timely, and its broad application is highly expected.

### Comparison to other methods

There have been various models to evaluate microbiome responses. Early *in vitro* gut microbiome models were based on large-scale bioreactors that are low-throughput and due to the large volume of culturing, very costly owing to the considerable amount of compounds added. More recent advances in modeling the gut ecosystem includes realizing culturing of complex human gut microbiome in anaerobic intestine-on-a-chip models, enabling the observation of host-microbiome interactions[34]. However, for the purpose of high-throughput compound screening, these models are not easily adaptable. Our method has the advantage of easy set-up and is cost- and time-efficient to scale up. In terms of the metaproteomic sample analysis stage, a most recent study reported the development of a high-throughput stool metaproteomics workflow[35]. The protocol used Protifi S-trap to clean up the proteins in combination with TMT labeling and automation, which also greatly saves time. However, only around 5,000 microbial proteins were identified from a total of 290 human stool samples[35]. We will show in the following **Anticipated results** section that protein identifications are remarkably higher using our optimized protocol. Metaproteomic profiling derived from cultured samples of four human gut microbiomes resulted in a total of over 10,000 quantified protein groups with an average of over 5,000 per sample. This is comparable to previous studies using manual, label-free metaproteomics protocols[10, 36].

### Overview of the RapidAIM 2.0 protocol

As illustrated in **Figure 1**, the RapidAIM 2.0 protocol is divided into six sequential stages: a) microbiome collection, culturing and compound treatment; b) microbial cell washing; c) protein extraction and purification; d) protein digestion and desalting; e) TMT labeling and desalting; f) LC-MS/MS analysis and metaproteomic data analysis. First, individual fecal samples are cultured with or without compounds/stimuli of interest (**Figure 1a**). An anaerobic chamber with a 37 °C incubator capable of accommodating an orbital shaker is used. A 96-well liquid handler is recommended for liquid handling.

**Figure 1.**
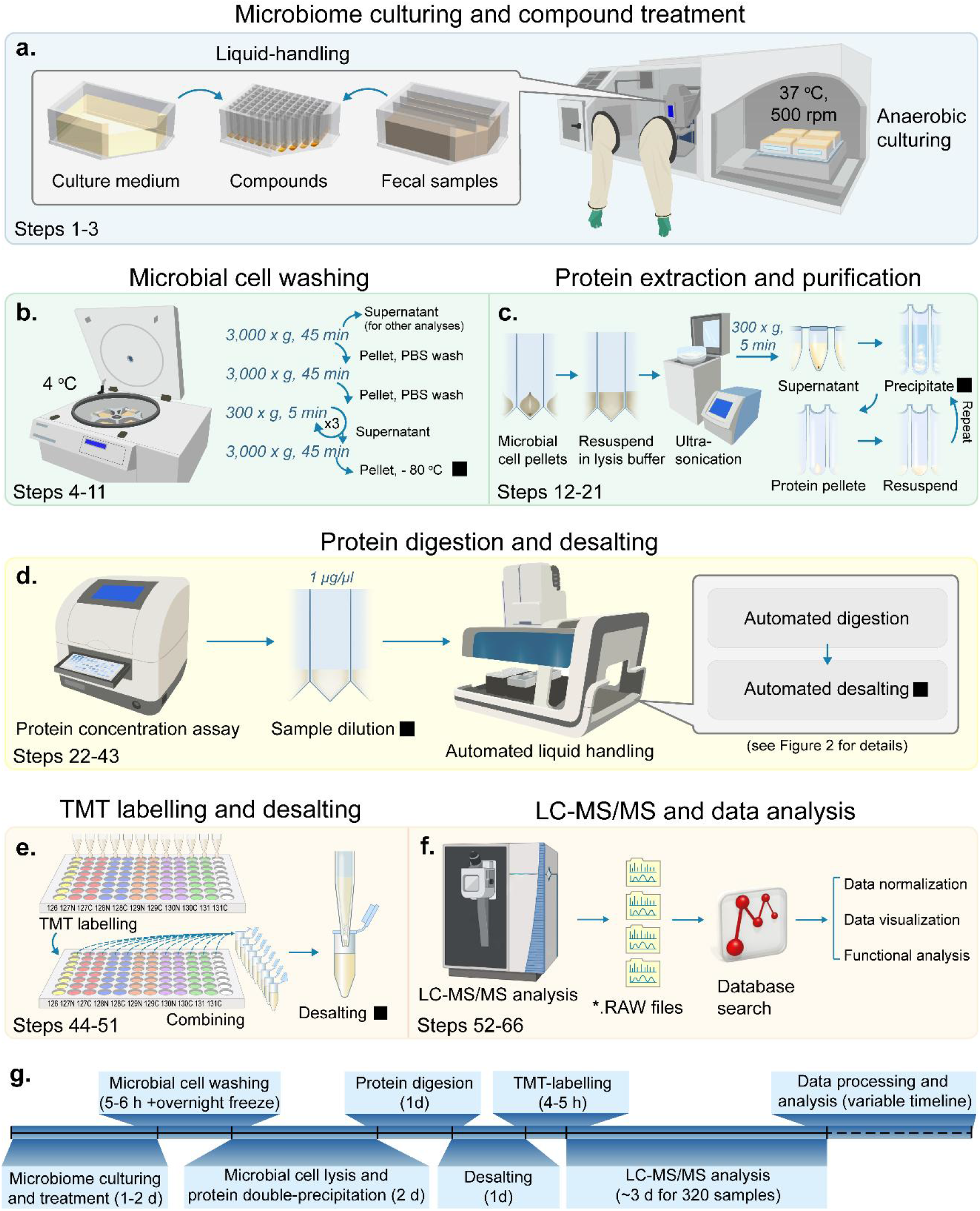
Overview of the RapidAIM 2.0 protocol workflow. **a.** Using an anaerobic chamber with a 37 °C incubator, individual fecal samples are cultured in the optimized medium with or without the stimuli of interest. A 96-well liquid handler is recommended for liquid handling. Sample plates are shaken on an orbital shaker at 500 rpm. **b.** Cultured microbiome samples are then washed with PBS buffer using a centrifuge with a deepwell plate rotor. **c.** Microbial cells are lysed in 96-well PCR plates using a cup-horn ultra-sonicator, proteins are then purified using a double-precipitation procedure. **d.** Proteins are then quantified and diluted, followed by automated digestion and desalting. **e.** Desalted peptides are then labeled with tandem mass tags (TMT) and each TMT11plex™ mix is then desalted again. **f.** Samples are finally analyzed with LC-MS/MS, and *.RAW files are subjected to database search and data analysis. g. Estimated timeline corresponding to an experiment of four 96-well plates (i.e. 320 samples). Filled squares indicate pause points.

Sample plates are shaken on an orbital shaker at 500 rpm for 18-24 hours. Either fresh human fecal samples or −80 °C stored biobank samples can be used for the culturing step. We previously showed that our culturing protocol maintains the functionality of individual microbiomes with both sample types [10, 23]. **Supplementary Methods 1 and 2** provide details for collecting and processing fresh stool samples, and collecting, processing and biobanking of the samples, respectively.

Next, microbial cells are purified using differential centrifugation then stored at −80 °C (**Figure 1b**). Proteins are then extracted from the cell lysate and purified using a double-precipitation procedure (**Figure 1c**). After protein quantification, samples are diluted to a recommended protein concentration of 1 μg/μl, followed by automated digestion and desalting procedures (**Figure 1d** and **Figure 2a-b**). For labs without automation capacity, 96-well plate-based manual digestion and desalting procedures can also be used (**Supplementary Methods 3**). However, we validated that the automation increased reproducibility compared to manual liquid handling (**Supplementary Figure S1**). Samples are then labeled using a TMT11plex™ reagents pre-aliquoted and dried into 96-well plates (**Figure 1e**). TMT labeling is quenched, pooled by rows and desalted prior to sample analysis using an LC-MS/MS. *.RAW files are subjected to database search using MetaLab software, and subsequent data analysis are preformed (**Figure 1f**). A timeline breakdown for the whole protocol is given by **Figure 1g**.

**Figure 2.**
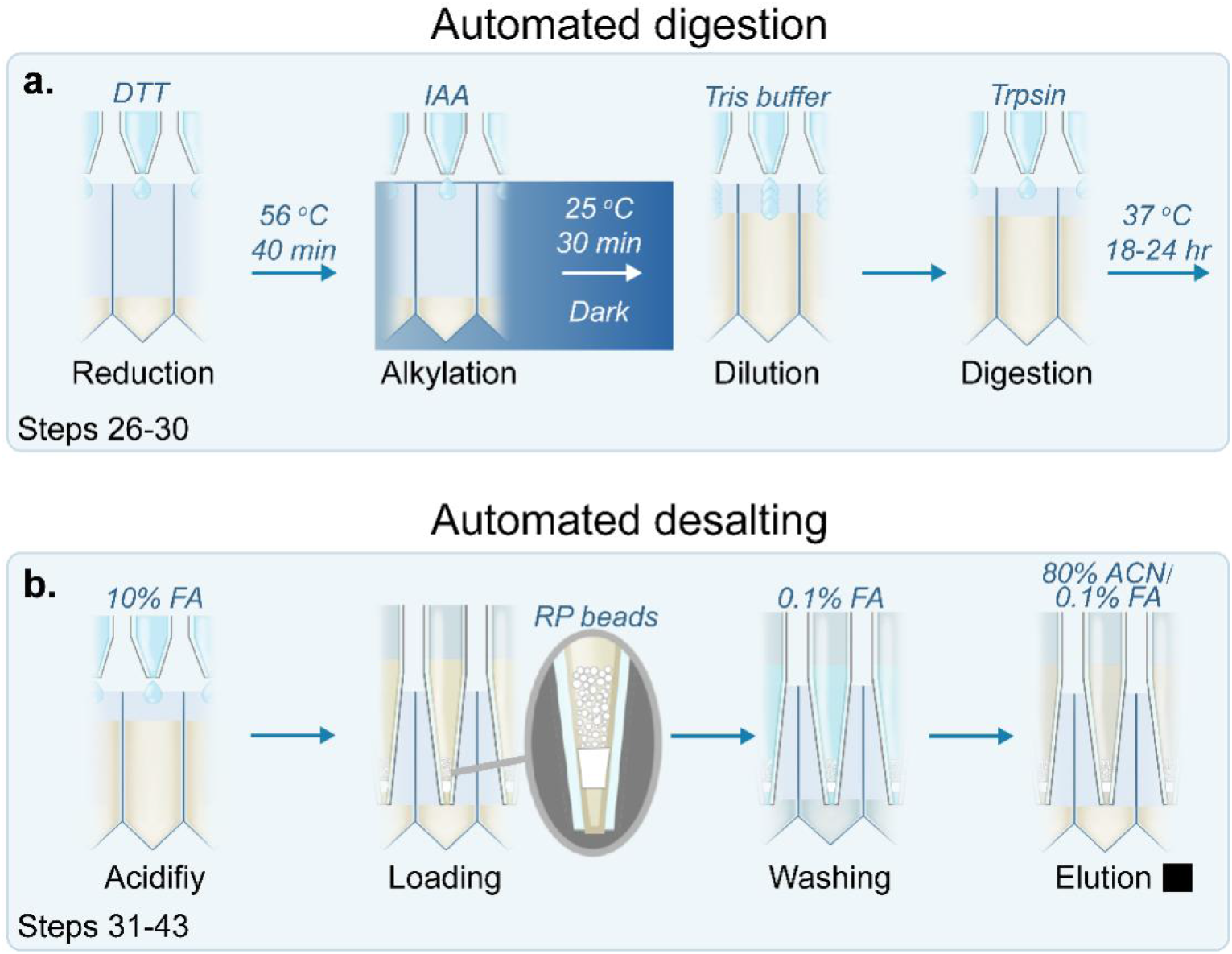
Automated protein digestion and desalting in 96-well plates. **a.** Protein samples are reduced with dithiothreitol (DTT) at 56 °C for 30 min, then alkylated with iodoacetamide (IAA) at 25 °C for 40 min in dark. After dilution with 100 mM Tris-HCl buffer (pH 8.0), protein samples are digested using trypsin at 37 °C for 18-24 hours on thermo-mixers. **b.** After digestion, samples are then acidified to pH 2-3 using 10% formic acid (FA, v/v), then loaded to pre-activated columns containing reverse phase (RP) beads. After being washed with 0.1% FA (v/v), tryptic peptides are eluted with 80% acetonitrile (v/v)/0.1% FA (v/v). Filled square indicates the pause point.

### Experimental design

1. **Plate layout.** The experimental design will be performed based on a 96-well format. Taking into consideration of the use of TMT11plex™, we recommend that an 8 lines x 10 columns plate layout is used for each 96-well plate. The first column will later on be used for the TMT reference sample, which will be generated after the desalting step. The last column will be left blank throughout the experiment.
2. **Randomization.** Compound treatments across all assay plates should be randomized. We provide the “96-well plate randomizer” tool in our iMetaLab Suite[37]s to assist researchers with the study randomization (https://shiny1.imetalab.ca/96_well_randomizer/). Randomizing within and across sample plates will be helpful to detect batch effects between plates, if any, and meet the criteria to apply batch removal tools[38]. Samples should be randomized again prior to LC-MS/MS analysis.
3. **Controls.** Vehicle controls which are microbiomes cultured in the absence of compound treatment but presence of compound vehicle should be included. For example, when compounds are pre-dissolved in DMSO, then the same amount of DMSO should be used in the vehicle control. Positive controls of known effect on the *ex vivo* human gut microbiome should also be included, such as FOS or kestose.
4. **Biological and technical replications.** An adequate number of biological replicates should be included and the sample size should be determined through power analysis[39]. In terms of technical replicates, for smaller scale studies or individualized studies, we recommend that each condition is carried out with technical triplicates. For large scale studies having sufficient power of biological replicates, the minimum requirement is that technical triplicates of negative and positive controls should be performed.
5. **Quality controls for LC-MS/MS.** For large-scale assays requiring a relatively long LC-MS/MS running time (e.g. > 2 days), quality control (QC) samples are necessary to ensure quality, reproducibility and comparability of results across different batches. We strongly suggest that the QC sample should be study-specific. For this, a mixture of a small aliquot from each TMT-labeled sample is recommended. It is also recommended that the researchers run the QC samples on LC-MS/MS to confirm sample quality, before labeling by TMT.

### Expertise needed to implement the protocol

This protocol requires that researchers are familiar with aseptic techniques in the microbiology lab to prevent the contamination of media, culture and work area, and to safely handle any biohazard materials. The protocol require access to an LC-MS/MS system, and the involvement of an experienced mass spectrometrist to ensure the instrument is performing at an optimal level for TMT analysis. For the bioinformatics workflow, the MetaLab software is designed to be easily used by any user through a graphical user interface. However, fundamental knowledge about metaproteomics database search workflow is required to ensure adequate setting of all search parameters. For data analysis, knowledge in multivariable statistical analysis and preliminary programming experience in R language are desired.

### Limitations

This protocol includes a metaproteomics analysis workflow following the gut microbiome culturing. As metaproteomics provides information on protein compositions, it does not include other information such as genomic and metabolite compositions. We recommend that in order to enable multi-omics analysis and other biochemical analysis of the samples, aliquots of samples should be saved at certain steps for such purposes. For example, the microbial cell-free supernatant of the microbial cell washing step may be taken/stored for possible metabolite analysis. At the final step of microbial cell washing, before pelleting the microbial cells, samples may be divided into two aliquots for metaproteomics and metagenomics, respectively. In addition, this protocol was developed for the aim of compound screening purpose which does not directly enable deep metaproteomics analysis. We recommend that the researchers save aliquots of protein lysate or digest for each sample and do fractionation-based deep metaproteomics with specific samples of interest following the first-pass screening analysis.

## Materials

### Biological materials

Fresh stool samples, or stool samples processed following our biobanking protocol and stored at −80 °C. See **Supplementary Methods 1** and **2** for stool sample collection and processing.

- CAUTION: Human feces is suggested to be handled as a level 2 biohazard (risk group 2), because they can contain pathogens such as bacteria, viruses, and parasites. Immunizations for those handling these samples may be available against some pathogens such as hepatitis. Personal protective equipment includes googles, gloves and lab coat should be worn when handling feces. Biosafety cabinets and centrifuges with lids to contain aerosols and spills should be used.

### Reagents

#### Microbiome culturing

- Potassium phosphate monobasic (Millipore-Sigma - Sigma-Aldrich, cat. no. P5655)
- Potassium phosphate dibasic (Millipore-Sigma - Supelco, cat. no. PX1570-1)
- Sodium chloride (Calbiochem - OmniPur®, cat. no. 7710)
- Magnesium sulfate heptahydrate (Millipore-Sigma - Sigma-Aldrich, cat. no. 230391)
- Calcium chloride, anhydrous (Millipore-Sigma - Sigma-Aldrich, cat. no. C5670)
  - CAUTION: Calcium chloride causes serious eye irritation, wear eye protection/ face protection.
- Tween® 80 (Millipore-Sigma - Sigma-Aldrich, cat. no. P4780)
- Sodium cholate hydrate (Millipore-Sigma - Sigma-Aldrich, cat. no. C9282)
- Sodium chenodeoxycholate (Millipore-Sigma - Sigma-Aldrich, cat. no. C8261)
- Peptone water (Millipore-Sigma - Sigma-Aldrich, cat. no. 70179)
- Bacto™ yeast extract (Becton, Dickinson and Company, cat. no. 212750)
- Sodium bicarbonate (Millipore-Sigma - Supelco, cat. no. SX0320)
- L-cysteine (Millipore-Sigma - Sigma-Aldrich, cat. no. C7352)
- 1-Kestose (TCI America, cat. no. K0032)
- Hydrochloric acid (HCl, Fisher Chemical, cat. no. A144S)
  - CAUTION: HCl causes severe skin burns and eye damage and may cause respiratory irritation. Avoid breathing dust/ fume/ gas/ mist/ vapors/ spray. Use only under a chemical fume hood. Use personal protective equipment.
- Hemin (Millipore-Sigma - Sigma-Aldrich, cat. no. 51280)
- Vitamin K1 ((Millipore-Sigma - Sigma-Aldrich, cat. no. V3501)

#### Metaproteomic sample processing

- Phosphate buffered saline, 10X solution (Fisher BioReagents, cat. no. BP399-4)
- Urea (Millipore-Sigma - Sigma-Aldrich, cat. no. U5378)
- Tris (hydroxymethyl)aminomethane (Calbiochem - OmniPur®, cat. no. 9230)
- Hydrochloric acid (HCl, Fisher Chemical, cat. no. A144S)
  - CAUTION: same as above
- Sodium dodecyl sulfate (Millipore-Sigma - Sigma-Aldrich, cat. no. L3771)
  - CAUTION: Sodium dodecyl sulfate causes skin, eye and respiratory irritation. Use personal protective equipment.
- Acetone (Millipore-Sigma - Sigma-Aldrich, cat. no. 179124)
  - CAUTION: Acetone is highly flammable liquid and vapor, causes serious eye irritation, and may cause drowsiness or dizziness. Keep away from open flames, hot surfaces and sources of ignition. Use only under a chemical fume hood. Use personal protective equipment.
- Acetic acid, glacial (HAc, Fisher Chemical, cat. no. A38-212)
  - CAUTION: Flammable liquid and vapor. Use personal protective equipment. Keep away from open flames, hot surfaces and sources of ignition. Use only under a chemical fume hood.
- Acetonitrile (Millipore-Sigma - Sigma-Aldrich, cat. no. 34851)
  - CAUTION: Highly flammable and toxic. Keep away from open flames, hot surfaces and sources of ignition. Use only under a chemical fume hood. Use personal protective equipment.
- Ethyl alcohol, anhydrous (Commercial Alcohols, cat. no. P016EAAN)
  - CAUTION: Ethanol is highly flammable. Keep away from open flames, hot surfaces and sources of ignition.
- Formic acid (Millipore-Sigma - Sigma-Aldrich, cat. no. F0507)
  - CAUTION: Flammable liquid and vapor, causes severe skin burns and eye damage and toxic if inhaled. Keep away from open flames, hot surfaces and sources of ignition. Use only under a chemical fume hood. Use personal protective equipment.
- cOmplete™ protease inhibitor cocktail (Millipore-Sigma - Roche, cat. no. 04693116001)
- Dithiothreitol (Millipore-Sigma - Sigma-Aldrich, cat. no. 43815)
- Iodoacetamide (Millipore-Sigma - Sigma-Aldrich, cat. no. I1149)
- Trypsin (Worthington Biochemical, cat. no. L5003740)
- DC Protein Assay Reagents A, B and S (Bio-Rad Laboratories, cat. no. 5000113, 5000114 and 5000115)
  - CAUTION: Reagent A causes severe skin burns and eye damage.

#### TMT labeling

- TMT10plex Isobaric Label Reagent Set plus TMT11-131C Label Reagent (Thermo Scientific, cat. no. A34808)
- 50% hydroxylamine for TMT experiments (Thermo Scientific, cat. no. 90115)
  - CAUTION: Causes skin irritation and serious eye damage. Use personal protective equipment.
- 1M triethylammonium bicarbonate (TEAB) for TMT experiments (Thermo Scientific, cat. no. 90114)
- Pierce™ quantitative colorimetric peptide assay (Thermo Scientific, cat. no. 23275)

### Equipment

- CRITICAL: we have tested and selected optimal plates and lids for each of the steps to achieve optimal material performance and efficiency.
- Culture plate: Corning® 96 well Polypropylene Deep Well Plate (Sigma-Aldrich, cat. no. CLS3960)
- Culture plate lid: Sealing mat for 2 mL square deep well plates (Sigma-Aldrich, cat. no. AXYAM2MLSQ)
- Lysis plate: 96-well non-skirted PCR plate (Thermo Scientific, cat. no. AB-0600)
- Lysis plate lid: Flat 8 cap strips (Thermo Scientific, cat. no. AB-0783)
- Precipitation plate: Corning® 96 well PP 1.2 mL cluster tubes (Sigma-Aldrich, cat. no. CLS4413)
- Precipitation plate lid: 96-well Polyethylene Cluster Tube 8-Cap Strips (Sigma- Aldrich, cat. no. CLS4418)
- Elution plate: 0.8ml 96-well storage plate (Thermo Scientific, cat. no. AB-0859)
- Elution plate lid: Nunc™ 96 Well Caps for 1.0mL Polystyrene DeepWell™ Plates (Thermo Scientific, cat. no. 278616)
- TMT plate: Nunc™ 96-Well Polypropylene Storage Microplates (Thermo Scientific, cat. no. 249944)
- TMT plate lid: 96 well cap natural (Thermo Scientific, cat. no. 276002)
- Reservoir: Axygen™ Single Well High Profile Reagent Reservoir (Axygen, cat. no. RESSW96HP)
- Centrifuge with a deepwell-plate rotor (Eppendorf, Model 5810R, cat. no. 0226270405810R; or equivalent)
- Ultra-sonicator with microplate horn and chiller (QSonica, cat. no. Q700MPXC)
- ThermoMixers with plate adaptors (Eppendorf, model ThermoMixer C – cat. no. 5382000023 with SmartBlock FP – cat. no. 5306000006 and ThermoTop Heated Cover – cat. no. 5308000003; or equivalent)
- Microplate reader (BioTek, model Synergy LX, cat no. SLXFATS; or equivalent)
- 96-channel electronic pipette (Eppendorf, epMotion® 96, cat. no. 5069000209; or equivalent)
- Automated 96-channel liquid handling platform (Hamilton, model Nimbus 96, cat. no. OPP041219; or equivalent)
- Desalting tip columns (e.g. IMCS, 04T-H6R05-1-10-96, 04T-H6R52-1-10-8 or equivalent)
- Vacuum concentrator and plate rotor (eg. Labconco Centrivap vacuum concentrator - cat. no. LCN 7310022 and plate rotor – cat. no. 7461900)
- LC-MS/MS system (e.g. an UltiMate 3000 RSLCnano system – cat. no. ULTIM3000RSLCNANO coupled with an Orbitrap Exploris 480 mass spectrometer – cat. no. BRE725533, Thermo Fisher Scientific)

### Software

- MetaLab (version 2.3.0, available via iMetaLab Suite [37] at https://imetalab.ca/)
- R (e.g. version 4.0.4, R Foundation)

### Reagent setup

CRITICAL: For a RapidAIM compound screening experiment designed with large sample sizes, the workflow can consume a lot of reagents compared to common microbiome culturing and metaproteomics analysis. To minimize lab cost, and ensure consistency across runs, we recommend the preparation of following stock solutions.

#### Culture medium stock solutions

Prepare stock solutions of the culture medium following details described in Table 1.

**Table 1:**
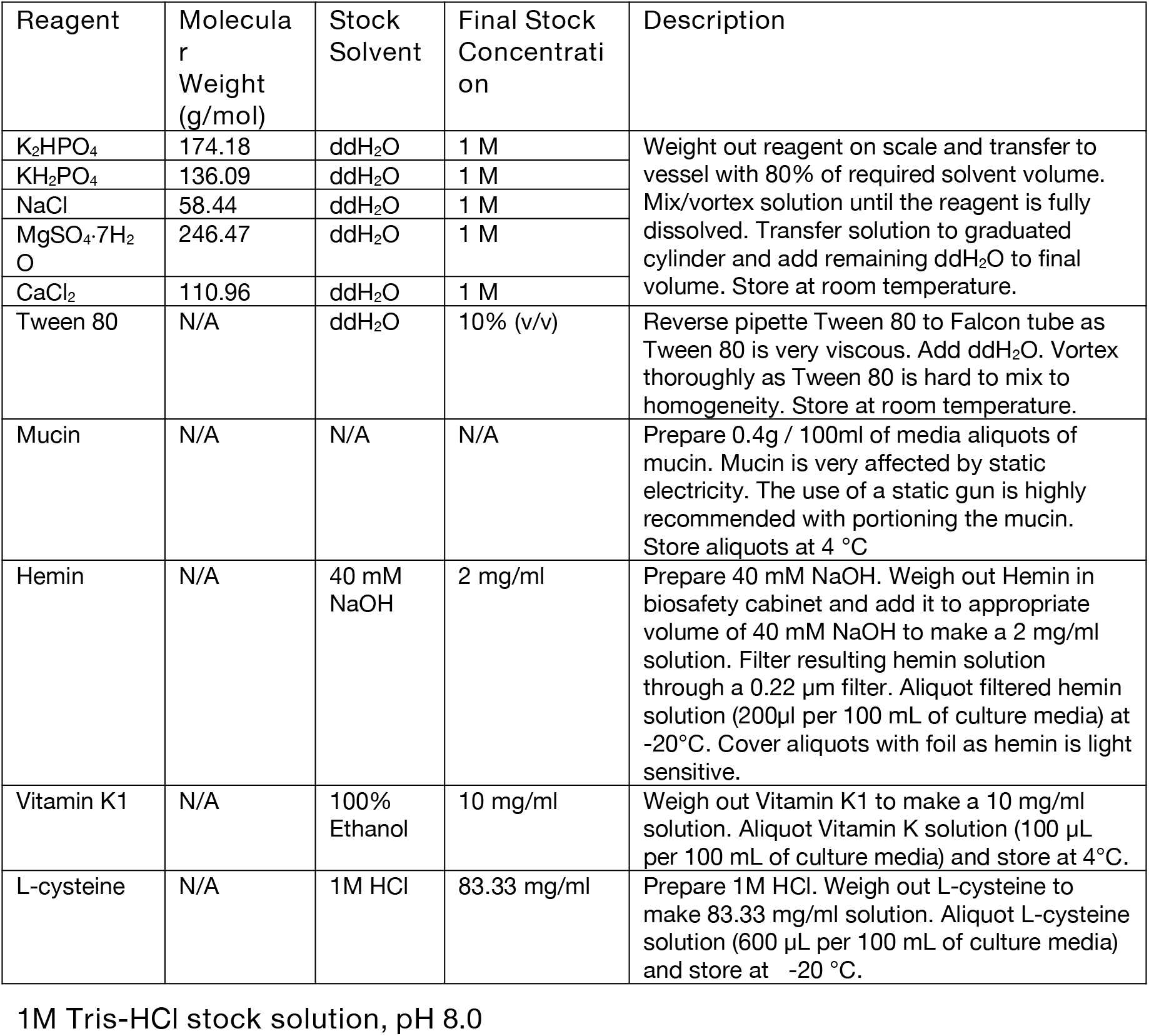
Preparation of stock solutions for RapidAIM culture media

Weigh out 12.11 g Tris base and add 80 mL of ddH_2_O. While mixing on a magnetic stirrer, observe pH and slowly add HCl solution to reduce the pH to 8.0. Top up the solution to 100 mL using ddH_2_O and double check pH.

#### TMT aliquot stock plates

Equilibrate the TMT11plex™ reagents (5 mg per channel) to room temperature, then add 300 μL of anhydrous ACN to each tube of the reagents, allow to dissolve for 5 minutes with occasional vortexing. Transfer each of the reagents to 15 mL Falcon tubes, and add 4700μL anhydrous ACN to each of the 15 mL tubes. Mix thoroughly. Aliquot the TMT reagents to 96 well plates (50 μL per well), each 11plex should be arranged in order in a row, as shown in **Figure 1e**. Freeze-dry at 4 °C and store at −80 °C.

#### Culture medium preparation

- CRITICAL: Culture media preparation begins with the mixing of autoclavable solutions and reagents the day before culturing

Prepare culture media consisting of the following composition: 2.0 g L^-1^ peptone water, 2.0 g/L yeast extract, 0.5 g/L L-cysteine hydrochloride, 2 mL/L Tween 80, 5 mg/L hemin, 10 μL/L vitamin K1, 1.0 g/L NaCl, 0.4 g/L K_2_HPO_4_, 0.4 g/L KH_2_PO_4_, 0.1 g/L MgSO_4_·H_2_O, 0.1 g/L CaCl_2_·2H_2_O, 4.0 g/L NaHCO_3_, 4.0 g/L porcine gastric mucin, 0.25 g/L sodium cholate and 0.25 g/L sodium chenodeoxycholate. Follow detailed procedures in Table 2 for medium preparation.

**Table 2.** Culture Media Preparation Table

| Step 1. Adding stock solution reagents into 80 mL ddH<sub>2</sub>O (final volume of 100 mL media): |  |  |
| --- | --- | --- |
| Reagent | Volume per 100 mL media | Description |
| K<sub>2</sub>HPO<sub>4</sub> stock | 258.3 µL | Pipette volumes of reagent stock solutions to autoclavable culture media bottle with 80% ddH<sub>2</sub>O of desired media volume. Put magnet stir bar inside bottle and mix on moderate speed. |
| KH<sub>2</sub>PO<sub>4</sub> stock | 330.6 µL |  |
| NaCl stock | 1540 µL |  |
| MgSO<sub>4</sub>·7H<sub>2</sub>O stock | 36.5 µL |  |
| CaCl<sub>2</sub> stock | 81 µL |  |
| Tween 80 stock | 2000 µL |  |
| Step 2. Weighing/adding dry reagents for culture media. |  |  |
| Reagent | Weight per 100 mL media | Description |
| Sodium cholate | 0.025 g | Weigh out each reagent using weigh paper and scale. Transfer reagent to same autoclavable culture media bottle. Rinse stock reagent powder on weigh |
| Sodium chenodeoxycholate | 0.025 g |  |
| Peptone water | 0.20 g |  |
| Yeast extract | 0.20 g | paper with some ddH<sub>2</sub>O into the bottle. Leave solution to mix for a total of 10 minutes. |
| NaHCO<sub>3</sub> | 0.40 g |  |
| Step 3. Transfers culture media to graduated cylinder and add remaining ddH<sub>2</sub>O to reach desired volume. Transfer back culture media to autoclavable bottle and have media autoclaved. |  |  |
| Step 4. Place culture media bottle in anaerobic chamber with lid ajar to deoxygenate overnight. |  |  |
| Step 5. Add non-autoclavable reagents to culture media: |  |  |
| Reagent | Volume or weight per 100 mL media | Description |
| Hemin stock | 200 µL | Sterilize and bring in appropriate pipettes and tips into anaerobic chamber. Pipette reagents into deoxygenated media. |
| Vitamin K1 stock | 100 µL |  |
| L-cysteine stock | 600 µL |  |
| Mucin | 0.40 g | Transfer 70% of media volume to Mucin aliquot in anaerobic chamber. Vortex media and mucin thoroughly until most of mucin is dissolved. Transfer back the mucin solution to remaining media solution and gently swirl until media is well mixed. |
| Step 6. Aliquot some of culture media into tube and measure dissolved oxygen content of media. Ensure oxygen content of media is less than 1% or 0.5 mg/mL |  |  |
| Step 7. Using previously measured aliquot, measure the pH of the culture media. Ensure culture media has a pH between ~7.0-7.4. |  |  |

#### Microbial cell lysis buffer

Make cell lysis buffer containing 8 M urea, 4% SDS in 100 mM Tris-HCl (pH = 8.0). For every 50 mL lysis buffer, add one Roche cOmplete™ tablet and sonicate to dissolve. Or use a cOmplete™ mini tablet for every 10 mL of lysis buffer.

- CRITICAL: Lysis buffer must be freshly prepared before use.

#### Protein precipitation solution

Prepare precipitation solution of 50%:50%:0.1% (v/v/v) of acetone: ethanol: acetic acid solution and keep at −20 °C for minimum overnight before use.

#### Protein resuspension buffer

6 M urea in 100 mM Tris-HCl, (pH 8)

#### 0.1 M DTT solution

Weigh 77 mg of DTT powder and add 5 mL ddH_2_O. Prepare freshly before use or prepare in advance and store solution at −80 °C.

#### 0.2 M IAA solution

Weigh 185 mg of IAA powder and add 5 mL ddH_2_O. Prepare freshly before use or prepare in advance and store solution at −80 °C.

- CRITICAL: Avoid light

#### Trypsin solution

100 mM Tris-HCl buffer containing 2 μg/mL trypsin, 1 mL is required per sample plate

- CRITICAL: Trypsin solution must be freshly prepared before use

#### Desalting buffers

- Wash buffer: 0.1% (v/v) FA in water
- Elution buffer: 0.1% FA in 80% ACN: 80% (v/v) ACN and 0.1% (v/v) FA in water
- Acidifying buffer: 10% FA: 10% (v/v) FA in water; and 5% FA: 5% (v/v) FA in water

#### TMT labeling solutions

- 100 mM TEAB in 20% ACN. Mix 10% (v/v) 1M TEAB, 20% (v/v) 100% ACN and 70% (v/v) HPLC-grade water.
- 0.8% hydroxylamine in 100 mM TEAB (quencher). Mix 10% (v/v) 1M TEAB with 90% (v/v) HPLC-grade water to prepare 100 mM TEAB water solution. Then mix 1.6% (v/v) 50% hydroxylamine with 98.4% (v/v) 100 mM TEAB water solution.
  - CRITICAL: TMT labelling solutions must be freshly prepared before use.

#### LC-MS/MS buffers

- Buffer A: 0.1% (v/v) FA in HPLC-grade water.
- Buffer B: 80% (v/v) ACN and 0.1% (v/v) FA in HPLC-grade water.

### Equipment setup

Here we use an UltiMate 3000 RSLCnano system coupled with an Orbitrap Exploris 480 mass spectrometer system; their setups are as shown in **Tables 3** and **4**, respectively. However, users should optimize and find their most suitable parameters. Other usable models include an Orbitrap Exploris 240 and Orbitrap Eclipse Tribrid as recommended by ThermoFisher Scientific.

**Table 3.** Setup of UltiMate 3000 RSLCnano system

| Time (min) | LC gradient (%B) | Flow rate (μL/min) |
| --- | --- | --- |
| 0 | 5.0 | 0.300 |
| 108 | 35.0 | 0.300 |
| 113 | 80.0 | 0.300 |
| 117 | 80.0 | 0.300 |
| 117.01 | 2.0 | 0.300 |
| 120 | Stop run |  |
| LC gradients |  |  |
| Solvent A | 0.1% FA |  |
| Solvent B | 80% ACN, 0.1% FA |  |
| Injection volume | 2 μL |  |

**Table 4.**
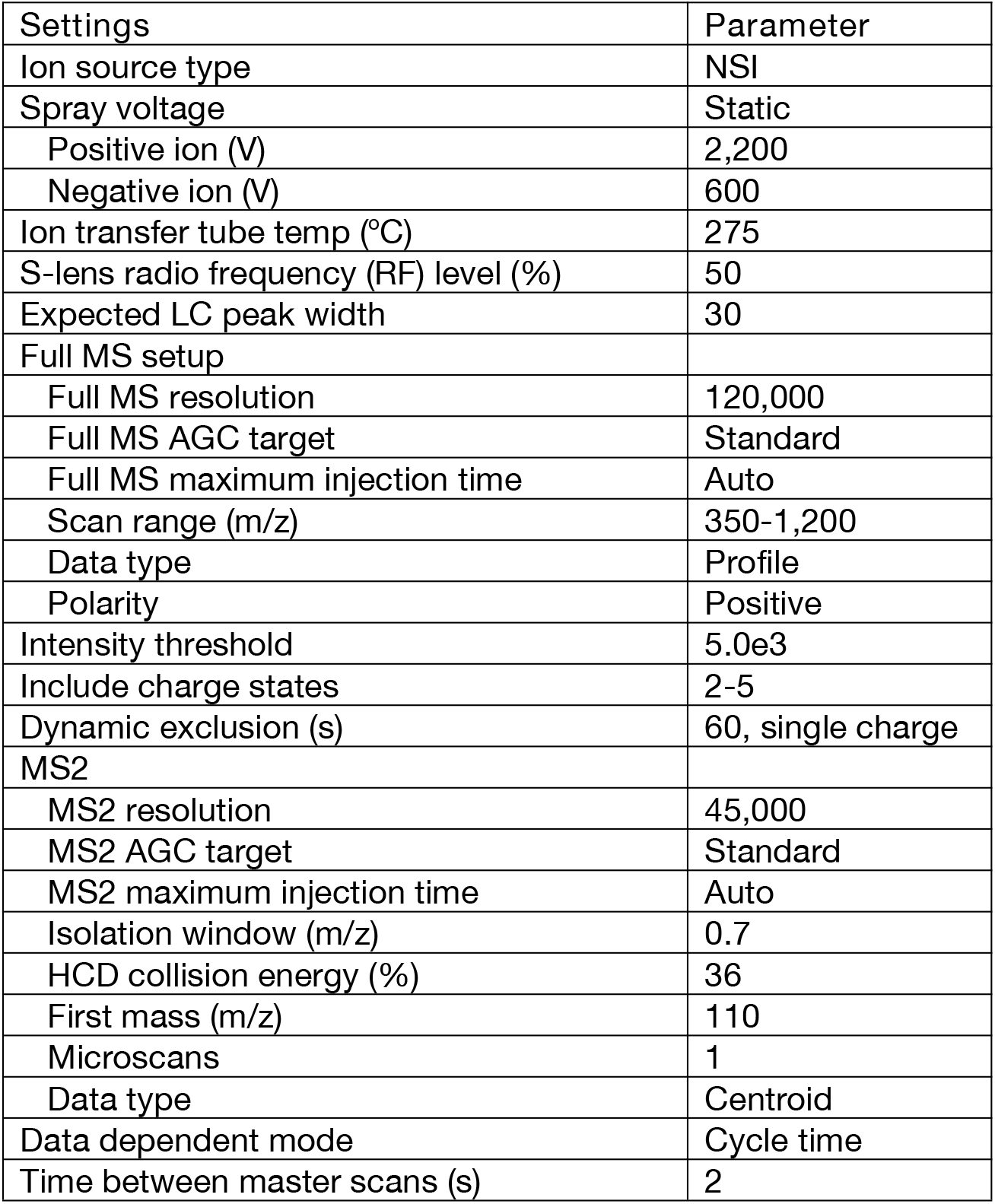
Setup of Orbitrap Exploris 480 mass spectrometer system

| Settings | Parameter |
| --- | --- |
| Ion source type | NSI |
| Spray voltage | Static |
| Positive ion (V) | 2,200 |
| Negative ion (V) | 600 |
| Ion transfer tube temp (°C) | 275 |
| S-lens radio frequency (RF) level (%) | 50 |
| Expected LC peak width | 30 |
| Full MS setup |  |
| Full MS resolution | 120,000 |
| Full MS AGC target | Standard |
| Full MS maximum injection time | Auto |
| Scan range (m/z) | 350-1,200 |
| Data type | Profile |
| Polarity | Positive |
| Intensity threshold | 5.0e3 |
| Include charge states | 2-5 |
| Dynamic exclusion (s) | 60, single charge |
| MS2 |  |
| MS2 resolution | 45,000 |
| MS2 AGC target | Standard |
| MS2 maximum injection time | Auto |
| Isolation window (m/z) | 0.7 |
| HCD collision energy (%) | 36 |
| First mass (m/z) | 110 |
| Microscans | 1 |
| Data type | Centroid |
| Data dependent mode | Cycle time |
| Time between master scans (s) | 2 |

#### Automated digestion deck set up

Set deck layout to have five plate/reservoir plate locations, two tip racks, a thermo block and a small-volume reservoirs location (for DTT and IAA). The plate locations are for four sample plates and one reservoir plate for trypsin-Tris-HCl solution. Two tip racks are for tips to be used in adding DTT and IAA, and trypsin-Tris-HCl solution, respectively.

#### Automated desalting deck set up

Set deck layout to have seven plate/reservoir locations and two tip racks. The plate locations are for one sample plate, one elution plate, two sample washing plates, and three reservoir plates for 100% ACN, 0.1% FA and 80% ACN + 0.1% FA solutions, respectively. The tip racks are for pipette tips and reverse-phase (RP) desalting columns, respectively.

## Procedure

Microbiome culturing and treatment [Timing 1-2 d]

1. Use a 96 channel liquid handler (e.g. epMotion® 96) or a multi-channel pipette to add 1 mL culture media and 100 μL stool samples into the square-well 96-deepwell plate containing compounds.
2. Mix sufficiently before covering the wells tightly with the perforated silicone gel mat and seal around using lab tapes to prevent popping up due to gas production in the culture. ? TROUBLESHOOTING: lids can pop up as a result of excess gas production in some individual microbiome. Make sure the silicone get mat is perforate at its thinnest place and use large gauge needles (e.g. gauge 18, L 1 ½ in., Millipore-Sigma - Sigma-Aldrich, cat. no. Z118044).Use lab tape to secure the lid when necessary.
3. Shake the deepwell plates at 500 rpm on an orbital shaker for 24-48 hours under 37 °C.
  - CRITICAL STEP: Anaerobic chamber must be maintained strictly anaerobic prior to and throughout the culturing process. Minimize the use of exchange chambers, and all materials to be used for the culturing should be pre-reduced at least over night to avoid surface-attached oxygen. Instead of using anaerobic chambers with open sleeves, we recommend the use of long-arm gloves.

Microbial cell washing [Timing 5-6 h]

4. After culturing, centrifuge culture plates at 3,000 g, 45 min, 4 °C.
5. Remove the microbial cell-free supernatant. The supernatant can be discarded or collected for pH, metabolome and/or exosome analysis.
6. Resuspend the pellets in 1 mL cold PBS buffer using a 96-channel liquid handler or a multi-channel pipette. If samples cannot be sufficiently resuspended using pipettes, firmly cover the plate with a silicon gel mat and vortex the plate at 2,000 rpm to mix well.
7. Centrifuge the plates again at 3,000 g, 45 min, 4 °C.
8. Add of 1 mL cold PBS buffer to resuspend the pellet again.
9. Centrifuge the plates at 300 g, 5 min, 4 °C to pellet debris. Carefully transfer the supernatant into a new deepwell plate.
10. Repeat step 6 for another two rounds, then centrifuge the plates at 3,000 g, 45 min, 4 °C.
11. Remove supernatant and store the plates at −80 °C before cell lysis.
  - CRITICAL STEP: During plate washing, the deepwell plates should be maximally sitting on ice and all liquid handling must be completed in a biosafety cabinet.
  - Pause point: Microbial cell pellets can be stored for ≤3 months at −80 °C.

Microbial cell lysis and protein double-precipitation [Timing 2 d]

Critical: The microbial cell lysis buffer contains SDS which can negatively affect the activity of trypsin during digestion. It is critical to purify the precipitated proteins from SDS. Here we use a double-precipitation workflow to make protein purification easy to handle for a 96-well plate format.

12. Prepare microbial cell lysis buffer (see Reagent setup) and precool to 8 °C.
13. Thaw microbial pellets in a 4 °C fridge.
14. Use a 96-channel liquid handler or a multichannel pipette to add 150 μL lysis buffer to the wells containing microbial cell pellets. Mix sufficiently to resuspend the pellets.
15. Carefully transfer the microbial cell suspensions to a 96-well PCR plate. Cover with plate lids.
  - CRITICAL STEP: Avoid foaming during transfer to the PCR plate. If there is an air gap at the bottom of the wells, centrifuge to pull all liquid down to the bottom. Having an air gap can affect sonication efficiency.
16. Sonicate with a cup-horn ultra-sonicator at 10 kHz, 10s-on and 10s-off cycle for 20 minutes (i.e. total sonication of 10 minutes).
  - CRITICAL STEP: Sonication produces heat that affects the sample; use a cooling system to maintain the sample’s surrounding temperature at 8 °C.
17. Centrifuge the sample plate at 300 g, 5 min to bring down droplets attached to the lids.
18. Transfer the cell lysate into a 1.2 mL 96-well cluster tube plate, and add 800 μL ice-cold protein precipitation solution. Cover with cluster lids and mix well. Precipitate at least overnight at –20 °C.
19. Centrifuge the sample plates at 3,000 g, 45 min, 4 °C. Carefully remove the supernatant. ? Troubleshooting: Crystal-like solids may be observed in some tubes as a result of urea crystallization. Slightly warming up the tubes with fingertip will help dissolve the urea.
20. Add 150 μL protein resuspension buffer to the pellets and mix at 2,000 rpm using a vortex mixer, until protein pellets in all wells are fully suspended.
21. Add 800 μL ice-cold protein precipitation solution. Cover with cluster lids and mix well. Precipitate at least overnight at –20 °C.
  - Pause point: protein precipitant can be stored for ≤1 months at −20 °C.

Protein concentration test and protein dilution [Timing 3-5 hours]

22. Centrifuge the sample plates at 3,000 g, 45 min, 4 °C and carefully remove the supernatant.
23. Add 100 μL protein resuspension buffer to the pellets and mix at 2,000 rpm using a vortex mixer, until protein pellets in all wells are fully suspended.
24. Determine protein concentration using the DC Protein Assay kit following the manufacturer’s instructions.
25. Dilute protein samples to reach a concentration of 1 μg/μL using the protein resuspension buffer. Aliquot 100 μL into 2 mL deepwell plates for protein digestion. The remaining can be kept as backup. Store at −20 °C before performing protein digestion.
  - Pause point: protein dilution can be stored for ≤1 months at −20 °C.

Protein digestion [Timing 1 d]

Critical: Protein digestion and desalting steps can be performed using an automated liquid handler equipped with a plate gripper and a thermos-block for plate heating and shaking. The digestion steps can also be performed using 96 channel liquid handlers or multi-channel pipettes (See Supplementary Material 3).

26. Add 10 μL 0.1 M DTT solution to each well. Incubate at 56 °C, 800 rpm for 30 minutes.
27. Cool the plates to room temperature.
28. Add 10 μL 0.2 M IAA solution to each well. Incubate at room temperature for 40 minutes.
  - CRITICAL STEP: IAA is light sensitive. Use black reagent reservoir for IAA and cover the 96-well plates during IAA incubation.
29. Add 1000 μL 100 mM Tris·HCl buffer containing 2 μg/mL trypsin (trypsin:proteins = 1:50).
  - CRITICAL STEP: The presence of 6M urea in the sample can significantly affect the activity of trypsin; this step dilutes the concentration of urea to 0.5 M. The trypsin solution must be freshly prepared. When using an automated liquid handler, a prompt dialog should be programmed to require users to add the trypsin-Tris·HCl buffer.
30. Mix sufficiently, cover the plates firmly and incubate at 37 °C, 800 rpm for 24 hours in ThermoMixers.

Desalting [Timing 1-2 hours]

31. After digestion, centrifuge at 300 g for 1 min to pull down any liquid condensate on the lid.
32. Acidify sample with 100 μL 10% FA and mix sufficiently. Use a pH strip to verify that the pH value is adjusted to 2-3.
33. Use an automated liquid handler and program to perform sample desalting steps. Set up deck layout as described in **Equipment setup**.
34. Add 800 μL 0.1% FA to each well of the two sample washing plates.
35. Condition the reverse-phase (RP) desalting columns by two cycles of up-and-down mixing in 100% ACN and two cycles of mixing in 0.1% FA. Recommended volume of mixing is 500-800 μL in each cycle. ? Troubleshooting: RP beads may float on liquid surface and attach to the side walls of the tips, increasing the mixing cycles and buffer volume will help to eliminate this issue.
36. Load samples to pre-activated reverse-phase (RP) columns by at least ten cycles of up-and-down mixing of 500-800 μL volume.
37. Wash the RP columns by two cycles of up-and-down mixing in the first sample washing plate, then move to the second sample washing plate and mix another two cycles.
38. Transfer 800 μL 80% ACN + 0.1% FA from the reservoir plate to the elution plate. ? Troubleshooting: 80% ACN + 0.1% FA solution can evaporate over time, resulting in a decrease of ACN concentration. Add a lid and remove by the automated gripper before liquid transfer, or use a prompt dialog to require users to add fresh buffer.
39. Elute samples in the RP columns by two cycles of 800 μL up-and-down mixing in the 80% ACN + 0.1% FA elution plate.
40. From the elution plate, aliquot 240 μL of the eluted solution to another 96-well plate to be used for TMT labeling.
41. From the elution plate, aliquot 40 μL of the eluted solution into a reservoir plate. Mix sufficiently. From the mixture, aliquot 240 μL to each well of the first column of the TMT labelling sample plates prepared in step 40.
  - Critical step: A mixture of all sample aliquots is recommended to be used as the reference sample for the TMT labeling.
  ? Troubleshooting: Volume of eluted sample can reduce over time since ACN evaporates quickly, therefore liquid handling and mixing should be performed as quickly as possible. To prevent evaporation during pipetting into TMT labeling sample plates, we suggest that the mixture is first transferred to a 15 mL Falcon tube(s).
42. The remainder can be kept as back-up or for label-free quantification in an LC-MS/MS.
43. Use a SpeedVac with a plate adapter to dry the samples under room temperature.
  - Pause point: Typically, dry peptides can be stored for ≤1 month at −20 °C or ≤6 months at −80 °C.

TMT-labeling and desalting [Timing 4-5 hours]

44. Take out TMT reagent plates and sample plates from freezers and naturally warm up to room temperature. Set ThermoMixers to 25 °C.
45. Add 20 μL 100 mM TEAB in 20% ACN solution to each sample well. Mix sufficiently using 600 rpm on an orbital shaker.
46. Aliquot 15 μL from each sample well to the corresponding wells of the TMT reagent plates. Cover with plate lids and incubate in the thermomixers at 25 °C, 600 rpm for 2 hours. Centrifuge plates at 300 g for 1 min to bring down any liquid on the lid.
47. Add 15 μL quencher (0.8% hydroxylamine in 100 mM TEAB) to each well. Cover with plate lids and react in the thermomixers at 25 °C, 600 rpm for 15 minutes. Centrifuge plates at 300 g for 1 min to bring down any liquid on the lid.
48. Acidify the samples by adding 60 μL 5% FA to each well. Mix sufficiently.
49. Combine each set of TMT11plex™ by taking 80 μL from each sample and combine all samples of a same row into a 96-deepwell plate.
50. Desalt by repeating steps 33-37.
51. Dry the desalted samples in a speedVac under room temperature. ? Troubleshooting: The selection of RP column material can affect the desalting of TMT samples. C18 beads should be used for this step.
  - Pause point: Typically, dry samples can be stored for ≤1 month at −20 °C before being analyzed in an LC-MS/MS.

LC-MS/MS analysis [Timing 2.5 hours per TMT11plex]

52. TMT quantitation is performed using a high resolution LC-MS/MS such as Orbitrap Exploris 480. An example of our LC-MS/MS parameters as a result of optimization is given in Equipment setup.
  - CRITICAL: High resolution MS/MS scanning is necessary for accurate ratio determination for TMT11plex™ experiments. The LC-MS/MS parameters should be optimized to achieve optimal performance including high quantitative accuracy/precision, coverage depth and minimizing co-ion interference and ratio compression.
53. Samples are resuspended at 1 μg/μL protein in 0.1% FA. Sufficiently mix using a vortex mixer then centrifuge at 14,000 g for 5 minutes before loading the supernatants to the LC-MS/MS sampler plate.
54. 1-2 μL of each sample is injected to the LC-MS/MS and analyzed following a 2-hour gradient.

Database search and data analysis [Timing variable timeline]

55. Database search of the LC-MS/MS rawfiles is performed using MetaLab 3.0. The software can be freely downloaded at http://imetalab.ca. It is recommended to run MetaLab 3.0 that is configured to use in a windows server.
56. Different workflows, e.g. MaxQuant or pFind workflows can be used for closed or open search, respectively. Here we demonstrate the use of MaxQuant for closed database search. The use of pFind follows a similar workflow.
57. Double-click MetaLab.exe to open the software.
58. Under the MaxQuant workflow option, select ‘Input data’ tab, click add to load the Rawfiles.
59. Under the ‘Database’ tab, click browse to load microbiome protein FASTA database (e.g. the IGC database, IGC.pep.fasta[40]).
60. Under the ‘Parameters’ tab, select ‘Carbamidomethyl (C)’ as fixed modifications, and ‘Acetyl (Protein N-term)’ and ‘Oxidation (M)’ as variable modifications.
61. Select ‘Trypsin’ as the enzyme and ‘Specific’ as the Digestion mode.
62. Select ‘Isobaric labelling’ as the quantification mode, and select TMT11plex™. In the bottom-right box, input the values of reporter ion isotopic distributions that are attached in the TMT reagent kit. CRITICAL STEP: Different batches of TMT11plex™ kit may have different values of reporter ion isotopic distributions. Make sure to keep record of lot number as well as the product sheet. Use the data corresponding to the correct lot number when performing the search.
63. Proceed to the ‘Settings’ tab. Select number of threads to use, ‘FTMS’ as the MS2 scan mode, and set up MaxQuant search parameters (here we use the default setting). Having a metadata setting is not mandatory, however labeling the samples by experimental groups will allow users to better visualize the data in the automated reports. CRITICAL STEP: It is recommended that sample-specific database and spectra clustering options are selected to optimize search speed. Please refer to Cheng et al.[41] about the spectral clustering method.
64. Proceed to the ‘Run’ tab, click ‘Start’ to begin the search. The console window will display the database search progress once started. ? Troubleshooting: The dialog in the ‘Run’ tab will help users to examine whether the current setting is correct for database search. A warning will be displayed and report the error for users to trouble shoot and correct the setting.
65. Once database search is done, a file folder named ‘maxQuant_search’ contains all result tables and an automate report for users to easily check the dataset quality.
66. Data pre-processing can be performed using the MSstatsTMT R package using proteinGroups.txt, evidence.txt tables and a user-customized msstatstmt_annotation file as the inputs. Please refer to the MSstatsTMT User Guide for detailed instructions. The processed data table can then be used for down-stream data analysis. ? Troubleshooting: Due to the multiplex nature of TMT11plex™, missing values usually appear across all eleven channels of a TMT sample. Imputation of the missing values may induce TMT sample-specific batch effects. Bioinformatic approaches that are compatible with missing values should be selected. For example, the nonlinear iterative partial least squares (NIPALS) algorithm is recommend use for principal component analysis (PCA).

## Trouble-shooting

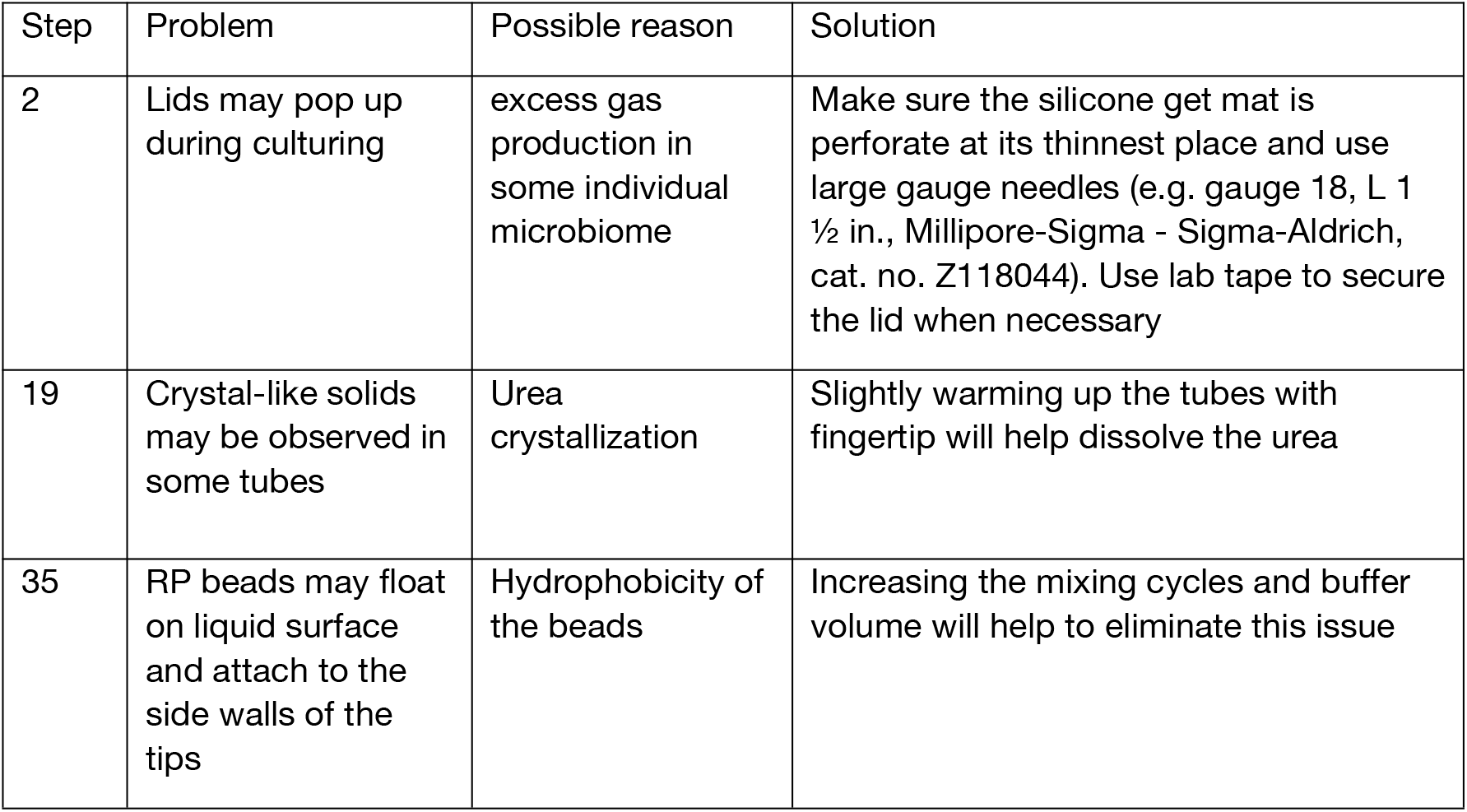
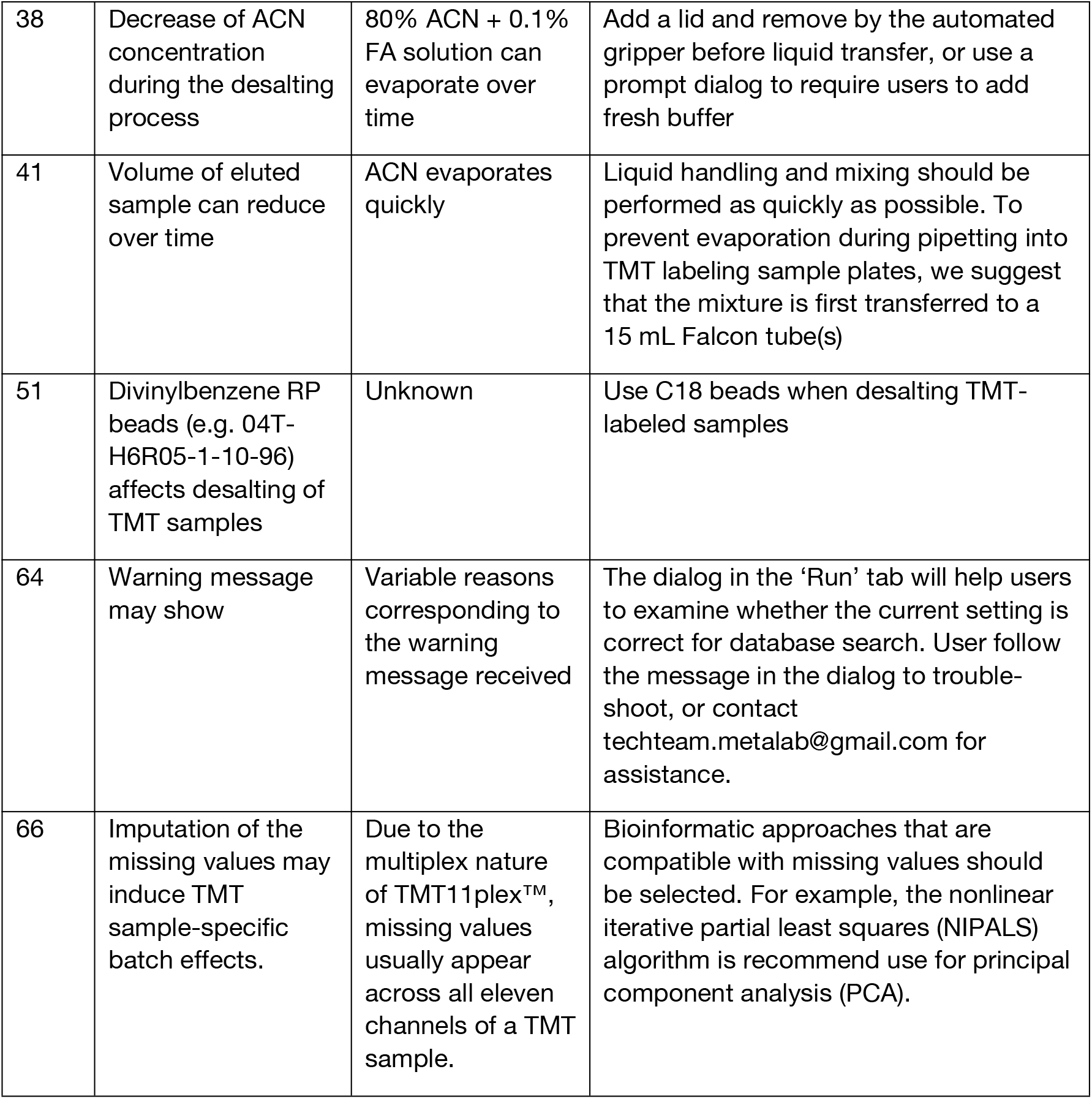

| Step | Problem | Possible reason | Solution |
| --- | --- | --- | --- |
| 2 | Lids may pop up during culturing | excess gas production in some individual microbiome | Make sure the silicone get mat is perforate at its thinnest place and use large gauge needles (e.g. gauge 18, L 1 ½ in., Millipore-Sigma - Sigma-Aldrich, cat. no. Z118044). Use lab tape to secure the lid when necessary |
| 19 | Crystal-like solids may be observed in some tubes | Urea crystallization | Slightly warming up the tubes with fingertip will help dissolve the urea |
| 35 | RP beads may float on liquid surface and attach to the side walls of the tips | Hydrophobicity of the beads | Increasing the mixing cycles and buffer volume will help to eliminate this issue |

## Timing

Steps 1-3, Microbiome culturing and treatment, 1-2 d

Steps 4-11, Microbial cell washing, 5-6 h

Steps 12-21, Microbial cell lysis and protein double-precipitation, 2 d

Steps 22-30, Protein digestion, 1 d

Steps 31-43, Desalting, 1-2 hours

Steps 44-51, TMT-labeling and desalting, 4-5 hours

Steps 52-54, LC-MS/MS analysis, 2.5 hours per TMT11plex

Steps 55-66, Database search and data analysis, variable timeline

## Anticipated results

RapidAIM was previously validated and applied for determining functional responses to a wide range of xenobiotic compounds and biotic components (see **Applications of the Method**). However, we have not showed its capacity when implemented with the 2.0 version protocol. Here we exemplify the described RapidAIM 2.0 workflow with demostration study, in which we evaluated the use of different human stool sample collection and processing workflows on individual gut microbiome responses to kestose, a prebiotic oligosaccharide. Four different stool sample processing strategies, namely gauze filtration (G), 100 μm vacuum filtration (V), 100 μm spin tube filtration (S) and 100 g spinning (100g) workflows were tested. Detailed protocols of these methods are described in **Supplementary Method S2**. A commercial kit GutAlive® was also evaluated following the manufacturer’s manual for its performance on live microbiome sample collection and preservation under room temperature for 24 hours and 72 hours before being processed using the 100 μm vacuum filtration protocol. All processed fecal samples were stored at −80 °C before performing the RapidAIM experiment. Microbiomes at 0 hour and 24 hours of culturing in the presence or absence of kestose were processed using the protocol described above. Samples of all comparisons were randomized before being labeled using TMT11plex™. An Ultimate 3000 RSLCnano system coupled to an Orbitrap Exploris 480 was used for the analysis of TMT-labeled samples using a two-hour gradient. Details of LC-MS/MS parameters are as described in Equipment setup. LC-MS/MS *.RAW files were searched against the IGC database [40] using MetaLab 2.3.0.

Altogether, 162 samples were labeled with a TMT11plex™ kit, resulting in 17 multiplexed samples. On average, 17,078 ± 24 MS/MS spectra were identified (MS/MS identification rate 24.2% ± 1.0%), resulting in 14,026 ± 570 identified peptides and 5,014 ± 142 protein groups per multiplexed sample (Mean ± SD, N = 17; **Figure 3a-c**). This is highly comparable to the previous label-free quantification in RapidAIM [10]. Dataset was then preprocessed using MSstatsTMT which includes a pipeline of spectrum-level normalization, protein summarization and protein-level normalization using the LogSum method[42]. Multivariate statistical analysis can then be performed using the processed dataset. Principal component analyses (PCA) show that all technical triplicates were well-clustered (**Figure 3d**). PCA and hierarchical clustering (**Figure 3d-e**) also show that for all the three tested individual microbiomes, samples were clustered by individual and condition (0 d baseline, vehicle, and kestose). No separation between the four different stool sample processing strategies were observed, suggesting that all four strategies were applicable for RapidAIM without impact on microbiome functional responses. Similarly, the test performed with the GutAlive kit using one individual microbiome sample showed clear separations by experimental conditions. While samples stored for 24 hours under room temperature in the GutAlive kit showed no separation with the samples that were processed on the day of collection, 72 hour sample showed a separation of the metaproteomics profiles from the other groups, indicating possible change of microbiome functionality that are specific to kestose uptake during the storage (**Supplementary Figure S2**).

**Figure 3.**
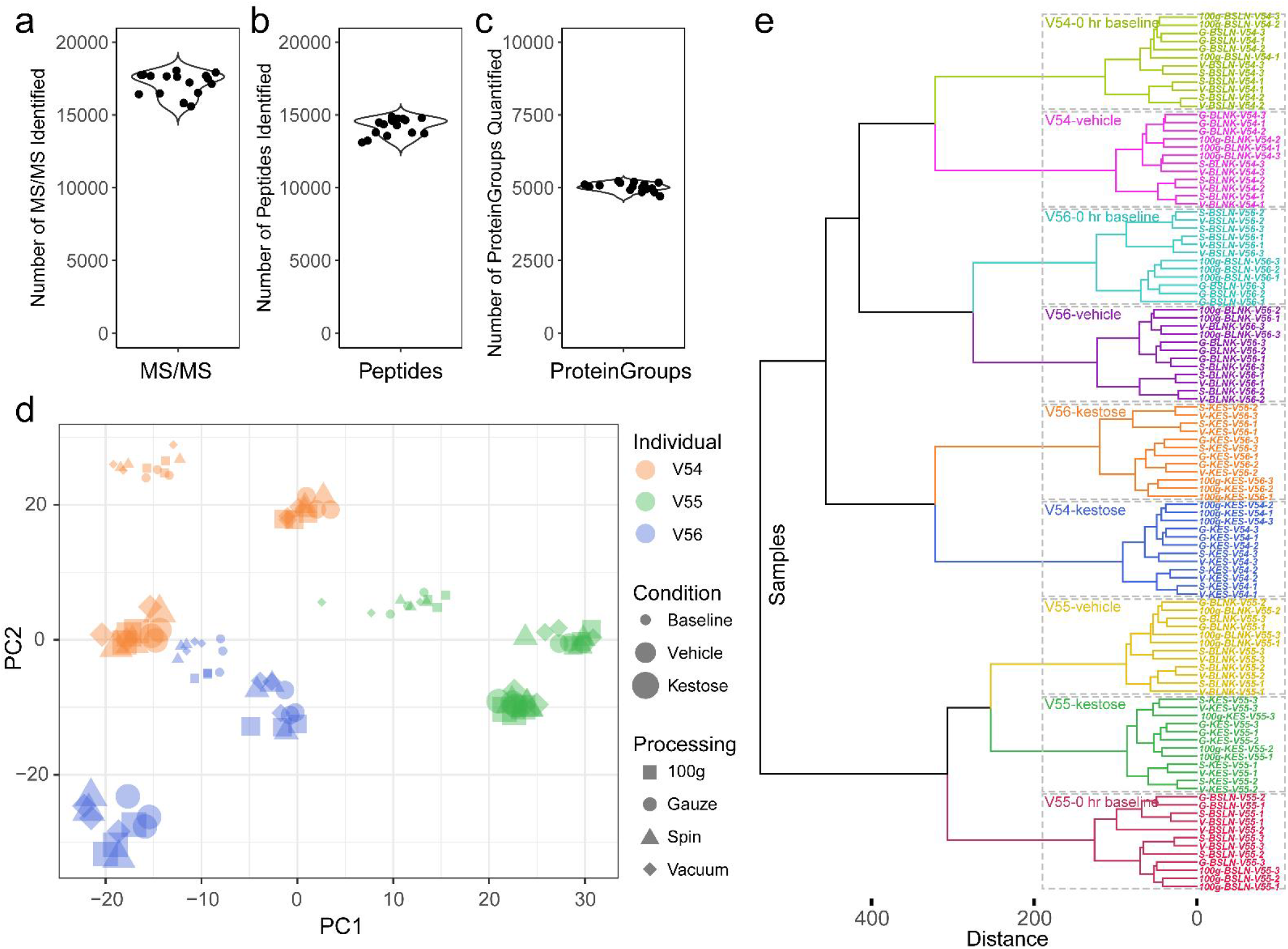
Evaluating effect of sample processing methods. a. Number of MS/MS identified in each sample. b. Number of peptides identified in each sample. c. Number of protein groups quantified in each sample. d. Principal component analysis of the sample processing study. Different colors indicate fecal samples from three different individuals. Sizes of data points represent different conditions (small - 0 hr baseline, medium – vehicle, large – baseline). Four different shapes indicate different processing methods. e. Hierarchical clustering of individual samples processed using different sample processing protocols and treated with or without kestose.

## Data availability

The mass spectrometry metaproteomics data have been deposited to the ProteomeXchange Consortium.

## Code availability

Source code of the Anticipated results section will be provided as an RNotebook file.

## Acknowledgements

This work was supported by the Government of Canada through Health Canada, Genome Canada and the Ontario Genomics Institute [OGI-156 and OGI-149], the Natural Sciences and Engineering Research Council of Canada [NSERC, grant no. 210034], and the Ontario Ministry of Economic Development and Innovation [ORF-DIG-14405 and project 13440].

## Notes

### Competing Interest Statement

D.F. co-founded MedBiome, clinical microbiomics company.

