## Supplementary_figures for "RapidAIM 2.0: a high-throughput assay to study functional response of human gut microbiome to xenobiotics"

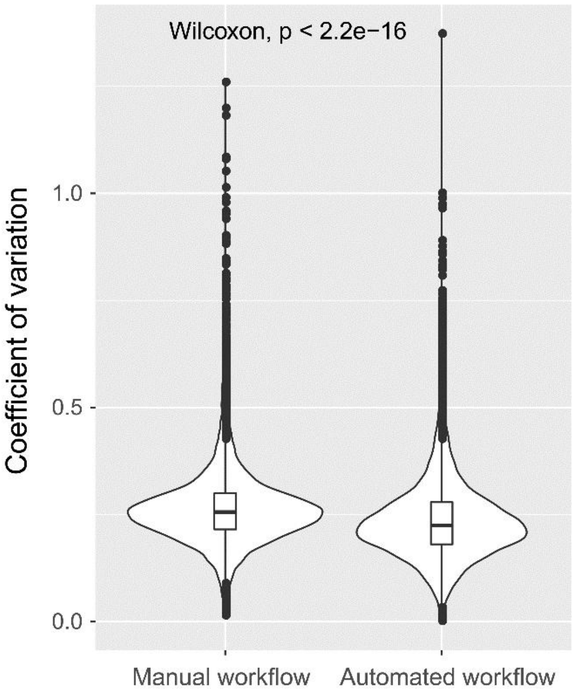


Supplementary Figure S1. Aliquots of a microbiome sample were processed using the manual workflow (as described in Supplementary Material 3) and the automated workflow (Steps 26-43 of the main text). For each method, technical triplicates were performed. Analysis of coefficient of variation based on peptide intensities showed a significant decrease using the automated workflow (Wilcoxon test).

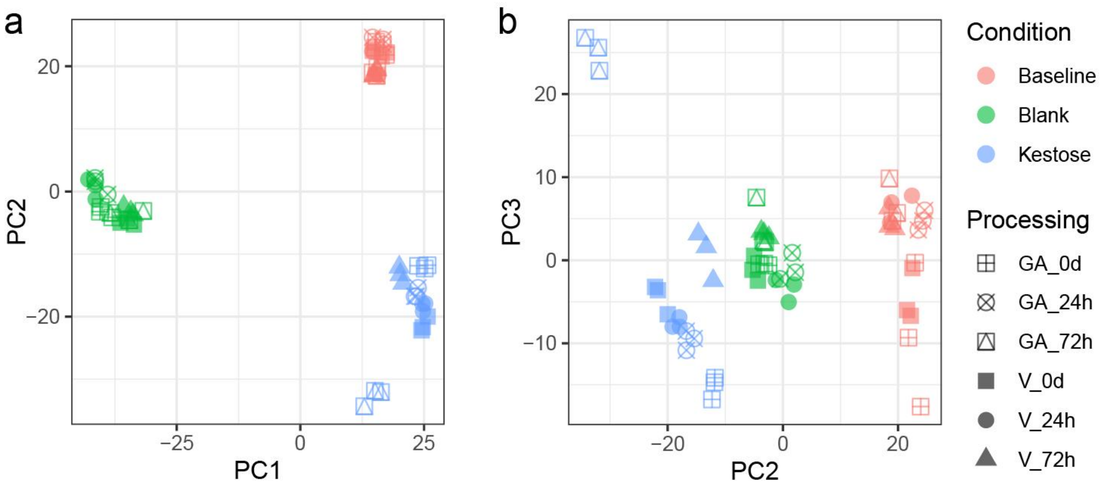


Supplementary Figure S2. Principal component analysis of the GutAlive kit assessment. Samples collected with the GutAlive kit and stored under room temperature for 24 hours and 72 hours were compared with samples collected with the collection kit described in Supplementary Methods 2 and stored at 4 °C for the same period of time for their maintenance of microbiome functionality by assessing functional responses to kestose using the RapidAIM protocol. a. PC1 vs PC2. b. PC2 vs PC3.
