## Supplementary_methods for "RapidAIM 2.0: a high-throughput assay to study functional response of human gut microbiome to xenobiotics"

Supplementary Method 1

### Fecal Sample Collection

- CAUTION: Institutional ethical approval must be obtained ensuring all samples are obtained with informed written consent and in accordance with relevant guidelines.
- CAUTION: Human feces is a level 2 biohazard (risk group 2) that can contain pathogens such as bacteria, viruses, and parasites. Immunizations for those handling these samples may be available against some pathogens such as hepatitis. Personal protective equipment includes googles, gloves and lab coat should be worn when handling feces. Biosafety cabinets and centrifuges with lids to contain aerosols and spills should be used. Samples, equipment, and waste materials need to be handled, decontaminated, and disposed of according to institutional biosafety guidelines.
- ADDITIONAL CONSIDERATIONS: De-identification of samples should be ensured by the study coordinator. Participants should be provided anonymity when provided kits for feces collection and for return of completed kits. Will your study require collection of dietary information and identification of feces consistency (eg; Bristol stool sample identification chart)?

### Fecal Sample Collection – On Site

### Materials and Reagents

Fecal sample collection kit – On site

- A flow chart of all kit components with instructions for completion
- Study specific documents (eg; 24-hr dietary recall and Bristol stool identification chart)
- Two leak-proof sterile return collection containers (eg; Thermo Scientific Samco Wide Mouth Bio-Tight Specimen Container, Fisher Scientific cat. no. 13-711-56)
    - Note: Container size should reflect the amount of feces needed for down-stream applications or storage. We mark a with a black to-fill line. Pre-weight dry collection container to calculate weight of feces following collection. Add label to be filled by participant with time and date of collection.

- Commode for feces collection (eg: Commode Specimen Collection System; Fisher Scientific cat. no. 02-544-208 or Sterile Collection Device, Therapak™; VWR cat. no. 76230-712)
- Disposable scoops for collection (eg; Bel-Art™ Sterile Sampler Spoons; Fisher Scientific cat. no. 14-429-D)
- Wooden tongue depressors (Fisher Scientific cat. no. S80332)
- Biohazard specimen transport bag (eg; Fisher Scientific cat. no. 22-310-094)
- Absorbent material for packing of biological liquids (eg; Therapak™ absorbent materials; Fisher Scientific cat. no. 22-130-041)
- Non-latex, chemically impermeable gloves
- Biohazard bag (eg; Bel-Art™ 1.5 mil thick biohazard disposal bag; Fisher Scientific cat. no. 03-411-700)
- Pens and markers for completing documents
- Transport box or bag (eg; Therapak Box; Fisher Scientific cat. no. 22-130-470)

Feces Stabilization Buffer – On site

- 1X phosphate buffered saline (pH 7.4; Wisent cat. no. 311-010-CL)
- Sodium thioglycolate (1 g/L; MilliporeSigma cat. no. 1066910500) or L-Cysteine (1mg/mL; MilliporeSigma cat. no. C7352)

### Equipment

- Serological pipettes and pipettor
- Autoclave or Disposable Filter Units (0.20 µm; eg; Fisher Scientific cat. no. FB12566504)
- Anerobic Chamber (eg; Sheldon Maunfacturing cat. no. BAA30022)
- Dissolved Oxygen Meter (eg; Extech Instruments cat. no. DO210)

### Procedure

Feces stabilization buffer preparation

1. Sterilize 1X PBS (pH 7.4) by autoclave or 0.20 µm vacuum filtering

2. Place into an anerobic chamber with cap loosened for >24-hr or until $O_2$-level is <0.5 mg/L
3. Sodium thioglycolate (1 g/L; MilliporeSigma cat. no. 1066910500) or L-Cysteine (1 mg/mL; MilliporeSigma cat. no. C7352)
4. Mix well.
5. Add 60 mL of deoxygenated feces stabilization buffer to leak-proof collection container. Ensure cap is tightened to prevent oxygenation of buffer.
    - NOTE: Feces stabilization buffer can be stored at 4°C. Before aliquoting ensure dissolved $O_2$-level is <0.5 mg/L. Weigh container pre- and post-buffer addition to note weight of buffer added during collection.

Feces Sample Kit Assembly and Collection

6. Assemble components listed above for sample collection kit.
7. Coordinator should add de-identifying number to the kit and deliver kit to participant pre-instructed on collection according to the following general flow-chart for sample collection instructions.
    a. Wash hands thoroughly. NOTE: Gloves can be worn.
    b. Empty bladder completely. NOTE: The feces samples should not be contaminated by urine.
    c. Collect stool sample into the disposable container. NOTE: The sample must not contact the toilet water.
    d. Using the scoop, collect feces from the middle of the specimen and directly put it into the dry collection container to the black fill line.
    e. Add 60 mL of deoxygenated feces stabilization buffer.
    f. Tightly close container.
    g. Label the container with the time and date the sample was collected.
    h. Place in biohazard specimen transport bag with absorbent material. Seal.
    i. Throw all wastes into the biohazard bag and seal.
    j. Place specimen bag and waste bag into secondary transport box or bag.
    k. Wash hands thoroughly.
    l. Store the sample at room temperature and return within 1 hour to the study coordinator. The kit will contain a de-identifying number.

m. Complete any study documents (eg; 24hr dietary recall form and Bristol stool identification page). Return to study coordinator.

Laboratory Sample Deliver and Processing

8. The coordinator should deliver fecal sample to the laboratory.
9. The fecal sample should be placed into the anerobic chamber immediately. NOTE: If sample cannot be processed immediately, sample can be placed at 4°C for up to 48-hr. Check that container lid is tight to ensure sample remains anaerobic.
10. Fecal samples should be processed to a 20% (w/v) slurry in sterile 1X PBS (pH 7.4) and 1 mg/mL L-Cysteine for immediate RapiAIM or to a 20% (w/v) slurry in sterile 1X PBS (pH 7.4) containing 10% (v/v) glycerol (MilliporeSigma cat. no. G5516) and 1 mg/ml L-Cysteine for aliquoting and storage at -80°C according to Method for Fecal Processing for RapidAIM and Storage in Supplementary Methods 2.

### Fecal Sample Collection – Off Site

- o CAUTION: Human feces is a level 2 biological reagent. Follow any regulations and requirements for proper packaging and shipment of level 2 biological materials.

### Materials and Reagents

- A flow chart of all kit components with instructions for completion
- Study specific documents (eg; 24-hr dietary recall and Bristol stool identification chart)
- Two leak-proof sterile return collection containers (eg; Thermo Scientific Samco Wide Mouth Bio-Tight Specimen Container, Fisher Scientific cat. no. 13-711-56)
    - o Note: Container size should reflect the amount of feces needed for down-stream applications or storage. We mark a with a black to-fill line. Pre-weight dry collection container to calculate weight of feces following collection. Add label to be filled by participant with time and date of collection.

- Commode for feces collection (eg: Commode Specimen Collection System; Fisher Scientific cat. no. 02-544-208 or Sterile Collection Device, Therapak™; VWR cat. no. 76230-712)
- Disposable scoops for collection (eg; Bel-Art™ Sterile Sampler Spoons; Fisher Scientific cat. no. 14-429-D)
- Wooden tongue depressors (Fisher Scientific cat. no. S80332)
- Biohazard specimen transport bag (eg; Fisher Scientific cat. no. 22-310-094)
- Absorbent material for packing of biological liquids (eg; Therapak™ absorbent materials; Fisher Scientific cat. no. 22-130-041)
- Non-latex, chemically impermeable gloves
- Pens and markers for completing documents
- Leak proof refrigerant shippers; nontoxic and food-grade gel refriderants (Fisher Scientific cat. no. 22-130-070). NOTE: We include 8 packs for large, insulated shipping box to maintain temperature at <8°C for up to 72-hrs.
- Two boxes to send sample kit components out to participants. One box to house feces stabilization buffer to be stored at 4°C by participant and one box to house cold packs to be stored at -20°C by participant.
- Bubble wrap for packing (Fisher Scientific cat. no. NC0513328
- Return tape (Find Tape.com LLC cat. no. JVCC RT 150/223 B)
- Insulated Transport Box for refrigerated return of level 2 biological substances (eg; Therapak™ Expanded Polystyrene Insulated Shippers; Fisher Scientific cat. no. 22-130-407
- Temperature recorder. NOTE: If temperature tracking is essential a recorder can be added to return package to monitor temperature throughout travel (eg; WarmMark Fisher Scientific cat. no. 22-111-029
- Shipping labels including exempt human specimen shipping label

Feces Stabilization Buffer – Off Site

- 1X phosphate buffered saline (pH 7.4; Wisent cat. no. 311-010-CL)
- Sodium thioglycolate (1g/L; MilliporeSigma cat. no. 1066910500) or L-Cysteine (1mg/mL; MilliporeSigma cat. no. C7352)
- Glycerol (10% (v/v); MilliporeSigma cat. no. G5516)

### Equipment

- Serological pipettes and pipettor
- Autoclave or Disposable Filter Units (0.20 µm; eg; Fisher Scientific cat. no. FB12566504)
- Anerobic Chamber (eg; Sheldon Maunfacturing cat. no. BAA30022)
- Dissolved Oxygen Meter (eg; Extech Instruments cat. no. DO210)

### Procedure

Feces stabilization and storage buffer preparation

1. Prepare 1X PBS (pH 7.4) + 10% (v/v) glycerol and sterilize by autoclave or 0.20 µm vacuum filtering
2. Place into an anerobic chamber with cap loosened for >24-hr or until $O_2$-level is <0.5 mg/l
3. Weigh L-Cysteine for addition to 1X PBS (pH7.4) + 10% (v/v) glycerol at 1 mg/mL.
4. Mix well.
5. Add 60 mL of deoxygenated feces stabilization buffer to leak-proof collection container. Ensure cap is tightened to prevent oxygenation of buffer.

- NOTE: Feces stabilization buffer can be stored at 4°C. Before aliquoting ensure dissolved $O_2$-level is <0.5 mg/L. Weigh container pre- and post- buffer addition to note weight of buffer added during collection. NOTE: For off-site collection we place a 2-week expiry date on these buffers and ask the participant request fresh buffer if sample not collected within two weeks of delivery.

Feces Sample Kit Assembly and Collection

6. Assemble components listed above for sample collection kit.
7. Coordinator should add de-identifying number to the kit and deliver kit to participant pre-instructed on collection according to the following general flow-chart for sample collection instructions.
   a. Upon receipt of kit, the participant is instructed to open the outer transport box. The participant should store feces stabilization buffer at 4°C in a refrigerator and box

containing cold packs in their freezer. Other components can remain at room temperature.

b. When participant is ready to collect feces sample, they should gather components of kit stored at room temperature, in refrigerator and in freezer.
c. Wash hands thoroughly. NOTE: Gloves can be worn.
d. Empty bladder completely. NOTE: The feces samples should not be contaminated by urine.
e. Collect stool sample into the disposable container. NOTE: The sample must not contact the toilet water.
f. Using the scoop, collect feces from the middle of the specimen and directly put it into the dry collection container to the black fill line.
g. Add 60 mL of deoxygenated feces stabilization buffer.
h. Tightly close container.
i. Label the container with the time and date the sample was collected.
j. Place specimen containers into secondary specimen bag with absorbent material and temperature recorder.
k. Throw all wastes into their garbage as appropriate.
l. Wash hands thoroughly.
m. Place two cold packs at bottom of insulated shipping box then place specimen bag into box and surround with remaining cold packs. The kit will contain a de-identifying number.
n. Close insulated box.
o. Complete any study documents (eg; 24hr dietary recall form and Bristol stool identification page) and place on top of insulated box.
p. Seal outer box with return tape and call coordinator to ship back to site. Study coordinator will have return shipping labels already on shipping box.

Laboratory Sample Deliver and Processing

8. The coordinator will receive shipment, remove any study documents and should deliver fecal sample to the laboratory.

9. The fecal sample should be placed into the anerobic chamber immediately. NOTE: If sample cannot be processed immediately, sample can be placed at 4°C for up to 48-hr from deposit. Check deposit time to calculate 48-hr deadline for processing.

11. Fecal samples should be processed to a 20% (w/v) slurry in sterile 1X PBS (pH 7.4) containing 10% (v/v) glycerol and 1 mg/ml L-Cysteine for aliquoting and storage at -80°C according to Method for Fecal Processing for RapidAIM and Storage in Supplementary Methods 2.

Supplementary Method 2

### Fecal Sample Processing for RapidAIM and for Long-term Storage at -80°C

- CAUTION: Human feces is a level 2 biohazard (risk group 2) that can contain pathogens such as bacteria, viruses, and parasites. Immunizations may be available against some pathogens such as hepatitis. Personal protective equipment includes googles, gloves and lab coat should be worn when handling feces. Biosafety cabinets and centrifuges with lids to contain aerosols and spills should be used. Samples, equipment, and waste materials need to be handled, decontaminated, and disposed of according to institutional biosafety guidelines.
- NOTE: If processing feces for immediate RapidAIM follow protocol below omitting the glycerol from the buffers which is required for long-term storage and viability of the isolated microbiome.

### Materials and Reagents

- Solid Glass beads 5mm (Fisher Scientific cat. no. 11-312C)
- 500 mL plastic bottles (eg: Nalgene™ Wide Mouth Packaging Bottles Leak Proof with Closure; Fisher Scientific cat. no. 03-313-14E
- 50 mL conical centrifuge tubes (Fisher Scientific cat. no. 14-959-49A)
- Wooden tongue depressors (Fisher Scientific cat. no. S80332)
- Gauze (Ultident cat. no. 400-4122)
- Funnels (Fisher Scientific cat. no. 10-500)
- Sterile Centifuge Filter Units 100 µm (eg; Steriflip™ Sterile Centrifuge Filter Units; MilliporeSigma cat. no. SCNY00100)
- 1X phosphate buffered saline (pH 7.4; Wisent cat. no. 311-010-CL)
- Sodium thioglycolate (1g/L; MilliporeSigma cat. no. 1066910500) or L-Cysteine (1mg/mL; MilliporeSigma cat. no. C7352)
- Spin filtration units with 100 µm filters (Ciro Manufacturing Corporation, customized tubes)

### Equipment

- Clinical centrifuge with rotor for 50 mL conical centrifuge tubes and rotor buckets with sealed lids
- Serological pipettes and pipettor
- Autoclave or Disposable Filter Units (0.20 µm; eg; Fisher Scientific cat. no. FB12566504)
- Anerobic Chamber (eg; Sheldon Maunfacturing cat. no. BAA30022)
- Dissolved Oxygen Meter (eg; Extech Instruments cat. no. DO210)

### Procedure

Feces stabilization and storage buffer preparation

- NOTE: Ensure buffer preparation precedes sample delivery by at least 24-hrs to ensure ample time for deoxygenation of storage buffer. Longer times will be necessary depending upon volume of buffer preparation. If using stored-buffers ensure they remained deoxygenated.
- Make excess buffer based upon final feces storage at 20% (w/v) in storage buffer.

1. Prepare 1X PBS (pH 7.4) + 10% (v/v) glycerol and sterilize by autoclave or 0.20 µm vacuum filtering.
2. Place into an anerobic chamber with cap loosened for >24-hr or until $O_2$-level is <0.5 mg/l
3. Weigh L-Cysteine for addition to 1X PBS (pH7.4) + 10% (v/v) glycerol at 1 mg/mL.
4. Mix well.
    - NOTE: Feces stabilization and storage buffer can be stored at 4°C. Before aliquoting ensure dissolved $O_2$-level is <0.5 mg/L. Feces Sample Kit Assembly and Collection

Anaerobic chamber Preparation

5. Bring materials and equipment into anaerobic chamber, including
    a. 15 mL Solid Glass beads – sterilized by autoclaving
    b. 1 x 500 mL plastic bottles – sterile or sterilized by autoclaving
    c. 10X 50 mL conical centrifuge tubes – sterile or sterilized by autoclaving

d. 10- and 25- mL serological pipettes and pipettor
e. Wooden tongue depressors – sterilized by autoclaving
f. Gauze – sterilized by autoclaving (for Processing Method 1) OR
g. Sterile Centrifuge Filter Units 100 μm (for processing Method 2)

Determining addition of stabilization and storage buffer for final 20% (w/v) fecal slurry

6. Amount of additional stabilization and storage buffer to be added to participant sample
   a. Weight of feces = (weight of feces sample + buffer + container) – buffer weight – container weight
   b. Total buffer needed for 20% fecal slurry = Weight of feces/0.20
   c. Amount of additional buffer to add = total buffer needed – 60ml added by participant

Processing Feces to 20% (w/v) slurry in stabilization and storage buffer

7. Place fecal sample into anaerobic chamber.
   - NOTE: The oxygen level of the buffer + feces can be measured and recorded. We exclude samples from further processing if $O_2$-level is >1 mg/L for samples collected 8-hrs prior as exposure to higher $O_2$-levels for extended periods indicated leaky containers and can affect microbiome viability/composition. Our experience shows variable O2-levels from 0.2-5 mg/L for samples collected on site or >8hrs from collection time.
8. Mix sample and buffer by vigorous shaking in original container.
9. Pour sample into 500 mL container. Depending on consistency of fecal slurry, wooden tongue depressors can aid in transfer.
10. Add glass beads and additional stabilization and storage buffer to make 20% (w/v) fecal slurry. Mix vigorously. Wooden tongue depressors can aid in mixing.

Clarifying Fecal Slurry

- NOTE: We have developed four alternative methods of clarifying fecal slurry of debris while maintaining microbiome composition. Each has pros and cons. The user can decide which method best suits their lab based upon these.
- Method 1 – Gauze Filtration
  Pro – gauze is inexpensive; required no additional equipment

Con – more time consuming

- Method 2 - 100 μm vacuum filtration

  Pro – fast- time efficient; very reproducible

  Con- filters ~4$/sample; requires clinical centrifuge; filter can clog requiring transfer to additional filter

- Method 3 - 100 μm spin tube filtration

  Pro – fast- time efficient; very reproducible

  Con- filters ~4$/sample; requires clinical centrifuge; spin tube can clog requiring transfer to additional spin tube

- Method 4 – 100 g spin

  Pro – fast- time efficient; very reproducible

  Con- requires clinical centrifuge

Method 1 – Gauze Filtration

11a. Place funnel onto 500 mL container and line funnel with 4-layers of gauze
12a. Pour a portion of fecal slurry into funnel.
13a. Use wooden tongue depressor to press slurry through
14a. Repeat until all slurry is cleared through gauze; changing gauze as needed
15a. Aliquot gauze-filtered 20% (w/v) fecal slurry and store at -80°C.

Method 2 – 100 μm Vacuum Filtration

11b. Aliquot fecal slurry into 50 mL centrifuge tubes
12b. Centrifuge at 100 g for 5 minutes
13b. Pour or pipette upper slurry into vacuum centrifuge tubes avoiding lower debris pellet and any floating debris.
14b. Vacuum filter slurry through 100 μm filter. We use a manual vacuum pipettor.
15b. Aliquot 100 μm vacuum-filtered 20% (w/v) fecal slurry and store at -80°C.

Method 3 – 100 μm Spin Tube Filtration

11c. Aliquot 20 mL fecal slurry into spin tube containing a 100 μm filtering insert
12c. Centrifuge at 100 g for 5 minutes
13c. Aliquot 100 μm spin tube -filtered 20% (w/v) fecal slurry and store at -80°C.

Method 4 – 100 g Spinning

11d. Aliquot fecal slurry into 50 mL centrifuge tubes

12d. Centrifuge at 100 g for 5 minutes.

13d. Pipette upper slurry into a new 50 mL centrifuge tube. Note: If there is floating debris following 100g spin pipette mid slurry – below debris- into a new 50 mL centrifuge tube.

14d. Aliquot 100g spun 20% (w/v) fecal slurry and store at -80°C.

Supplementary Method 3

96-well Plate Based Manual Digestion and Desalting Workflow

### Materials and Reagents

- Urea (Millipore-Sigma - Sigma-Aldrich, cat. no. U5378)
- Tris (hydroxymethyl)aminomethane (Calbiochem - OmniPur®, cat. no. 9230)
- Hydrochloric acid (HCl, Fisher Chemical, cat. no. A144S)
    - CAUTION: same as above
- Sodium dodecyl sulfate (Millipore-Sigma - Sigma-Aldrich, cat. no. L3771)
    - CAUTION: Sodium dodecyl sulfate causes skin, eye and respiratory irritation. Use personal protective equipment.
- Acetone (Millipore-Sigma - Sigma-Aldrich, cat. no. 179124)
    - CAUTION: Acetone is highly flammable liquid and vapor, causes serious eye irritation, and may cause drowsiness or dizziness. Keep away from open flames, hot surfaces and sources of ignition. Use only under a chemical fume hood. Use personal protective equipment.
- Acetic acid, glacial (HAc, Fisher Chemical, cat. no. A38-212)
    - CAUTION: Flammable liquid and vapor. Use personal protective equipment. Keep away from open flames, hot surfaces and sources of ignition. Use only under a chemical fume hood.
- Acetonitrile (Millipore-Sigma - Sigma-Aldrich, cat. no. 34851)
    - CAUTION: Highly flammable and toxic. Keep away from open flames, hot surfaces and sources of ignition. Use only under a chemical fume hood. Use personal protective equipment.
- Ethyl alcohol, anhydrous (Commercial Alcohols, cat. no. P016EAAN)
    - CAUTION: Ethanol is highly flammable. Keep away from open flames, hot surfaces and sources of ignition.
- Formic acid (Millipore-Sigma - Sigma-Aldrich, cat. no. F0507)
    - CAUTION: Flammable liquid and vapor, causes severe skin burns and eye damage and toxic if inhaled. Keep away from open flames, hot surfaces and sources of ignition. Use only under a chemical fume hood. Use personal protective equipment.

- cOmplete™ protease inhibitor cocktail (Millipore-Sigma - Roche, cat. no. 04693116001)
- Dithiothreitol (Millipore-Sigma - Sigma-Aldrich, cat. no. 43815)
- Iodoacetamide (Millipore-Sigma - Sigma-Aldrich, cat. no. I1149)
- Trypsin (Worthington Biochemical, cat. no. L5003740)
- ReproSil-Pur 120 C 18-AQ, 10 µm (Dr. Maisch GmbH, cat. no. r10.aq.0010)
- DC Protein Assay Reagents A, B and S (Bio-Rad Laboratories, cat. no. 5000113, 5000114 and 5000115)
    - CAUTION: Reagent A causes severe skin burns and eye damage.

#### Equipment

- Precipitation plate: Corning® 96 well PP 1.2 mL cluster tubes (Sigma-Aldrich, cat. no. CLS4413)
- Precipitation plate lid: 96-well Polyethylene Cluster Tube 8-Cap Strips (Sigma-Aldrich, cat. no. CLS4418)
- Lid for self-packed desalting tips: Nunc™ 96 Well Caps for 1.0mL Polystyrene DeepWell™ Plates (Thermo Scientific, cat. no. 278616)
- Desalting plate: Axygen® 96-well Clear Round Bottom 2 mL Polypropylene Deep Well Plate (Axygen, cat. no. P-DW-20-C-S)
- Elution plate: 0.8ml 96-well storage plate (Thermo Scientific, cat. no. AB-0859)
- Elution plate lid: Nunc™ 96 Well Caps for 1.0mL Polystyrene DeepWell™ Plates (Thermo Scientific, cat. no. 278616)
- Reservoir: Axygen™ Single Well High Profile Reagent Reservoir (Axygen, cat. no. RESSW96HP)
- Four thermomixers each with a plate adaptor (Eppendorf, model ThermoMixer C – cat. no. 5382000023 with SmartBlock FP – cat. no. 5306000006 and ThermoTop Heated Cover – cat. no. 5308000003; or equivalent)
- 96-channel electronic pipette (Eppendorf, epMotion® 96, cat. no. 5069000209; or equivalent)
- 20 µL filtered tips (Vertex, cat. no. 4237NAF)
- ReproSil-Pur 120 C18-AQ, 10 µm (Dr. Maisch GmbH, cat. no. r10.aq.0010)
- Centrifuge with a deepwell-plate rotor (Eppendorf, Model 5810R, cat. no. 0226270405810R; or equivalent)

### Reagent setup

1M Tris-HCl stock solution, pH 8.0

Weigh out 12.11 g Tris base and add 80 mL of dd$H_2O$. While mixing on a magnetic stirrer, observe pH and slowly add HCl solution to reduce the pH to 8.0. Top up the solution to 100 mL using dd$H_2O$ and double check pH.

Protein resuspension buffer

6 M urea in 100 mM Tris-HCl, (pH 8)

0.25 M DTT solution

Weigh 193 mg of DTT powder and add 5 mL dd$H_2O$. Prepare freshly before use or prepare in advance and store solution at -80 °C.

0.5 M IAA solution

Weigh 462 mg of IAA powder and add 5 mL dd$H_2O$. Prepare freshly before use or prepare in advance and store solution at -80 °C.

- CRITICAL: Avoid light

Trypsin solution

100 mM Tris-HCl buffer containing 2 µg/mL trypsin, 1 mL is required per sample

- CRITICAL: Trypsin solution must be freshly prepared before use

Desalting buffers

- Wash buffer: 0.1% (v/v) FA in water
- Elution buffer: 0.1% FA in 80% ACN: 80% (v/v) ACN and 0.1% (v/v) FA in water
- Acidifying buffer: 10% FA: 10% (v/v) FA in water; and 5% FA: 5% (v/v) FA in water

Self-packed desalting tip columns

- Slice crosses into each cap of the Nunc™ 96 Well Caps
- Cut the 20 µL filtered tips to the lower end of the filtering frits
- Insert the cut 20 µL filtered tips into the opening of the well caps
- Place the filter-tip-inserted well caps on a desalting plate
- For each set of 96 well desalting tips, weigh 600 mg of ReproSil-Pur 120 C18-AQ, 10 µm beads and add 6000 µL of 100% ACN.

- Immediately after sufficiently mixing to resuspend the C18 beads, use a 96-channel liquid handler or a multi-channel pipette to aliquot 50 µL of the resuspension into each desalting tip.
  - CRITICAL: C18 beads sediments quickly, mix again it happens.
- Centrifuge at 50-100 g for 1 minutes to remove ACN.

### Procedure

#### Digestion [Timing 1 d]

1. Use a 96 channel liquid handlers or multi-channel pipettes to add 4 µL 0.25 M DTT solution to each well. Incubate at 56 °C, 800 rpm for 30 minutes in a ThermoMixer.
2. Cool the plates to room temperature.
3. Add 4 µL 0.5 M IAA solution to each well. Incubate at room temperature for 40 minutes.
   - CRITICAL STEP: IAA is light sensitive. Use black reagent reservoir for IAA and cover the 96-well plates during IAA incubation.
4. Add 1000 µL 100 mM Tris·HCl buffer containing 2 µg/mL trypsin (trypsin:proteins = 1:50).
   - CRITICAL STEP: The presence of 6M urea in the sample can significantly affect the activity of trypsin; this step dilutes the concentration of urea to 0.5 M. The trypsin solution must be freshly prepared. When using an automated liquid handler, a prompt dialog should be programmed to require users to add the trypsin-Tris·HCl buffer.
5. Mix sufficiently, cover the plates firmly and incubate at 37 °C, 800 rpm for 24 hours in ThermoMixers.

#### Desalting [Timing 4-6 hours]

6. After digestion, centrifuge at 300 g for 1 min to pull down any liquid condensate on the lid.
7. Use a 96 channel liquid handlers or multi-channel pipettes to acidify sample with 100 µL 10% FA and mix sufficiently. Use a pH strip to verify that the pH value is adjusted to 2-3.
8. Activate the self-packed desalting columns by adding 300 µL of 100% ACN to each tip. Centrifuge at 100 g for 1 minutes.
9. Repeat step 8. Discard liquids in the desalting plate.
10. Equilibrate the tips by adding 300 µL of 0.1% FA. Centrifuge at 200 g for 2 minutes.
11. Repeat step 10. Discard liquids in the desalting plate.
12. Load 300 µL samples to the activated columns. Centrifuge at 200 g for 2 minutes.

? TROUBLESHOOTING: Due to difference of viscosity in the samples, some lysate may not be fully spun down, gradually increase the centrifugal force to bring all sampled through the filter.

CRITICAL: Do not let the beads go dry.

13. Repeat step 12 until all samples are loaded.
14. Wash the columns by adding 300 μL of 0.1% FA. Centrifuge at 200 g for 2 minutes.
15. Repeat step 14.
16. Replace the desalting plate into the elution plate.
17. Add 200 μL 80% ACN + 0.1% FA to each desalting tip, centrifuge at 100 g for 1 minute.
18. Repeat step 17.
19. From the elution plate, aliquot 120 μL of the eluted solution to another 96-well plate to be used for TMT labeling.
20. From the elution plate, aliquot 20 μL of the eluted solution into a reservoir plate. Mix sufficiently. From the mixture, aliquot 120 μL to each well of the first column of the TMT labelling sample plates prepared in step 19.
21. The remainder can be kept as back-up or for label-free quantification in an LC-MS/MS.
22. Use a SpeedVac with a plate adapter to dry the samples under room temperature.
    - Pause point: Typically, dry peptides can be stored for ≤1 month at -20 °C or ≤6 months at –80 °C.
